# Functional promoter polymorphisms govern the differential expression of HMG-CoA Reductase gene in rat models of essential hypertension

**DOI:** 10.1101/614669

**Authors:** Abrar A. Khan, Poovitha Sundar, Vinayak Gupta, Vikas Arige, S. Santosh Reddy, Madhu Dikshit, Manoj K. Barthwal, Nitish R. Mahapatra

## Abstract

3-Hydroxy-3-methyl glutaryl-coenzyme A reductase (*Hmgcr*) encoding the rate-limiting enzyme in the cholesterol biosynthesis pathway is a candidate gene for essential hypertension (EH). However, the regulation of *Hmgcr* in rat models of hypertension (viz. Spontaneously Hypertensive Rats [SHR] and its normotensive control Wistar/Kyoto [WKY] strain) is unknown. Here, we show that Hmgcr transcript and protein levels are diminished in liver tissues of SHR as compared to WKY. Consistently, a number of other rat models of hypertension display diminished cholesterol levels as compared to corresponding control strains. Sequencing of the *Hmgcr* promoter in SHR/WKY reveals three variations: A-405G, C-62T and a 11 bp insertion (-393_-382insTGCGGTCCTCC) in SHR. Moreover, SHR-*Hmgcr* promoter displays higher activity than WKY-*Hmgcr* promoter in various cell lines. Transient transfections of *Hmgcr*-promoter mutants and *in silico* analysis suggest altered binding of Runx3 and Srebf1 across A-405G and -393_-382insTGCGGTCCTCC sites. Indeed, chromatin immunoprecipitation assays confirm differential binding of Runx3/Srebf1 to *Hmgcr* promoter leading to diminished expression of *Hmgcr* in SHR as compared to WKY under basal/cholesterol-modulated conditions. Taken together, this study provides mechanistic insights for the altered *Hmgcr* expression in these models of EH, thereby unravelling the links of this gene to hypertension and associated cardiovascular disease states.

## INTRODUCTION

Essential hypertension, a major risk factor for cardiovascular diseases, is often associated with dysregulated cholesterol levels (1, 2). Cholesterol is the precursor of corticosteroids which play a crucial role in blood pressure (BP) regulation (3). Hence, 3-hydroxy-3-methylglutaryl-coenzyme A (HMG-CoA) reductase gene (human: *HMGCR*, mouse/rat: *Hmgcr*) that codes for a ∼ 97 kDa glycoprotein catalysing the rate-limiting step in the cholesterol biosynthesis pathway is a susceptibility gene for hypertension (4-6).

Transcriptome analysis in adrenal glands of two independent genetic rodent models of hypertension viz. spontaneously hypertensive rats (SHR) and blood pressure high (BPH) mice revealed ∼2.5 to 3-fold elevated *Hmgcr* expression as compared to their respective controls [viz. Wistar/Kyoto (WKY) rat and blood pressure low (BPL) mice] (7). *Hmgcr* was also over-expressed (by ∼2.6-fold) in the liver of BPL mice as compared to BPH. Interestingly, sequencing of the regulatory regions in the *Hmgcr* locus (viz. promoter, exons and flanking introns) from BPH, BPL and blood pressure normal (BPN) strains revealed multiple variations in the proximal promoter and coding regions. Two variations in the *Hmgcr* promoter domain (C-874T and C-470T) altered the binding affinities of transcription factors n-Myc, Max and c-Fos resulting in the differential expression of *Hmgcr* in these models of essential hypertension (8). However, the mechanism of regulation of *Hmgcr* expression in rat models of hypertension (viz. SHR and WKY) has not been explored so far.

SHR strain, the most widely used model for essential hypertension, was developed based on a breeding program where elevated BP was used for selection in Wistar rats (9, 10). The normotensive control strain for SHR, Wistar/Kyoto (WKY) strain, was also generated in the same breeding program by inbreeding of the original Wistar colony (from which the SHR strain originated) (11). SHR parallels human essential hypertension with co-morbidities (viz. insulin resistance, left ventricular hypertrophy, hypertriglyceridemia and abdominal obesity) (12-14). In the current study, we sequenced the proximal *Hmgcr* promoter in SHR as well as WKY and discovered three promoter variations [viz. A-405G, C-62T and a 11 bp insertion (-393_-382insTGCGGTCCTCC)]. Next, we assessed the functional impact of these variations by systematic computational and experimental analyses. Our results reveal that A-405G and - 393_-382insTGCGGTCCTCC altered the binding affinity of Runx3 and Srebf1 transcription factors and modulated the *Hmgcr* expression in these rat models of human essential hypertension. Interestingly, intracellular cholesterol level also regulates *Hmgcr* expression via Runx3 and Srebf1. HMGCR transcript level positively correlated with the transcript level of SREBF1 in various human tissues. In addition, this study for the first time to our knowledge, provides evidence of *Hmgcr* regulation by Runx3 under basal and cholesterol-modulated conditions.

## MATERIALS AND METHODS

### Blood pressure QTLs and LOD scores

The information about the blood pressure QTLs along with their LOD scores and all the genes harboured within a particular QTL was retrieved from the Rat Genome Database (15). Next, comparative genome analysis of the rat genomic regions with mouse and human genomic regions at the *Hmgcr* locus was performed using mVISTA (16). The mVISTA browser uses AVID alignment algorithm that employs global alignment for sequence comparison of the input queries.

### Rat strains and tissue samples

All animal procedures were approved by the Institutional Animal Ethics Committee at Indian Institute of Technology Madras and CSIR-Central Drug Research Institute, Lucknow, India. Male Spontaneously Hypertensive Rats (SHR) and Wistar/Kyoto (WKY) rats (n=6) were obtained from National Laboratory Animal Centre, CSIR-Central Drug Research Institute, Lucknow, India. These animals were housed in polypropylene cages under controlled room temperature at 25 °C on a 12 hr light-dark cycle and allowed to acclimatize for 2 weeks. All the animals received water and food *ad libitum*. After 5 weeks of a normal chow diet regimen, the animals were sacrificed and liver tissues from these animals were collected in RNAlater (Thermo Fisher, Waltham, USA).

### PCR amplification and molecular cloning of rat *Hmgcr* promoter

The genomic DNA isolated from SHR and WKY liver tissues was PCR-amplified by Phusion™ High-Fidelity DNA polymerase (Finnzyme, USA) using the following primers: forward, FP-RatHMGCR-1KbPro: 5’-CGG**GGTACC**GTGGGTAGGTATATCCGGGCT-3’ and reverse, RP-RatHMGCR-1KbPro: 5’-CCG**CTCGAG**AGCCACCTCGTGGAACCAGG-3’. *Kpn*I and *Xho*I sites were added to the forward and reverse primers, respectively which are underlined and indicated in bold. The purified PCR-amplified promoter fragments (∼1 kb) were cloned between *Kpn*I and *Xho*I sites in firefly luciferase reporter vector pGL3-Basic (Promega, USA). The resulting plasmids were named as SHR-*Hmgcr* and WKY-*Hmgcr* which contain −985 bp to +20 bp region of SHR and WKY *Hmgcr* promoter sequence, respectively. Here, the cap site indicates +1 position and the numberings of the nucleotide positions are with respect to the cap site. The resultant *Hmgcr*-promoter-reporter plasmids were confirmed by sequencing using RV3 (5’-CTAGCAAAATAGGCTGTCCC-3’) and GL2 (5’-CTTTATGTTTTTGGCGTCTTCCA-3’) primers.

### Sequencing and analysis of the *Hmgcr* promoters

The sequence of approximately ≃1 kb *Hmgcr* promoter domains from SHR and WKY rats was determined by Sanger sequencing of the promoter plasmids followed by multiple sequence alignment using ClustalW (17). The *Hmgcr* promoter regions harbouring polymorphisms were analyzed to predict putative transcription factor binding sites using online tools viz. ConSite (18), TFSEARCH (19), P-Match (20) and LASAGNA (21) at 65-95% cut-offs.

The *Hmgcr* promoter domains from different mammals harbouring the identified variations were mined from UCSC genome browser. The conservation of nucleotides in regions harbouring the polymorphisms across various mammals was aligned and analyzed using GeneDoc (22).

### Generation of site-directed mutations at SHR/WKY-*Hmgcr* promoter regions harbouring polymorphisms

In order to identify functional polymorphisms in the SHR/WKY Hmgcr promoter domains, mutations were generated at the regions of interest by site-directed mutagenesis using the SHR/WKY promoter construct as template and a primer pair with the desired mutation (Table 1). The resulting promoter mutants were named as S405, W394, SDM, S62 and W62 (Fig.4A). The authenticity of these mutations was confirmed by sequencing.

**Table 1.**
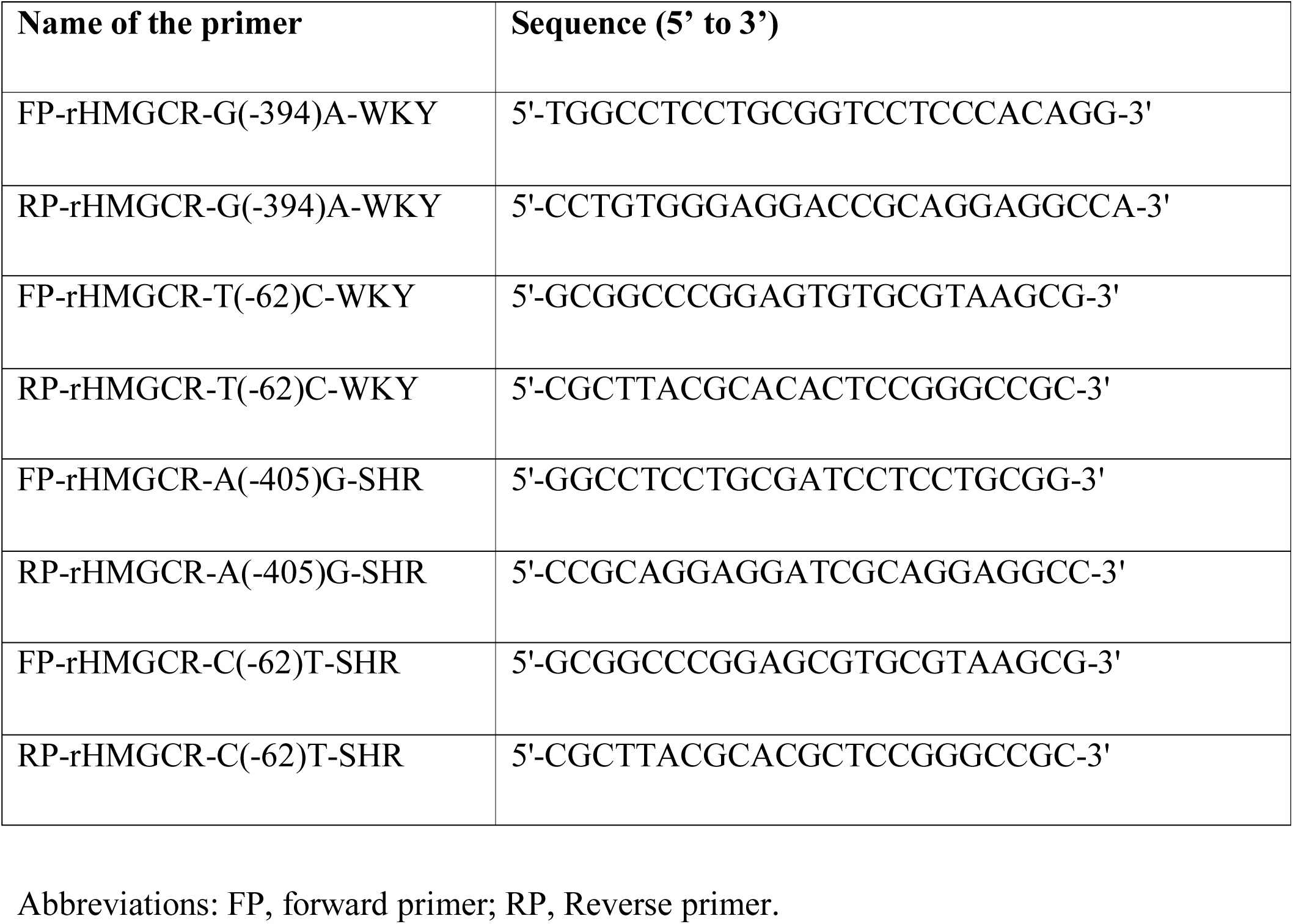
Primers used to generate Hmgcr promoter mutant constructs using site-directed mutagenesis.

### Tissue-specific expression of *HMGCR, SREBF1* and *RUNX3*

*HMGCR, RUNX3* and *SREBF1* expression data in various human tissues was mined from the EMBL-EBI expression atlas (http://www.ebi.ac.uk/gxa; Table 2) and GTEx portal (https://gtexportal.org/home/; Table S2). Those tissues which exhibited at least 10% *HMGCR* expression compared to that of the ovary with the highest TPM (82) and TPM cut-off of at least 1 for *SREBF1* and *RUNX3* were selected from the EMBL-EBI expression atlas for analysis. Likewise, tissue samples showing TPM of at least 1 for SREBF1 and RUNX3 were shortlisted from the GTEx portal.

**Table 2.** Transcript levels of HMGCR, RUNX3 and SREBF1 across various human tissues retrieved from EMBL-EBI expression atlas*.

| **No.** | **Tissue sample** | ***HMGCR***<br>**(TPM)** | ***RUNX3***<br>**(TPM)** | ***SREBF1***<br>**(TPM)** |
| --- | --- | --- | --- | --- |
| 1 | adipose tissue | 6 | 2 | 5 |
| 2 | adrenal gland | 17 | 39 | 23 |
| 3 | breast | 13 | 3 | 29 |
| 4 | colon | 11 | 1 | 4 |
| 5 | liver | 37 | 1 | 30 |
| 6 | lung | 37 | 24 | 13 |
| 7 | lymph node | 12 | 54 | 11 |
| 8 | ovary | 82 | 2 | 41 |
| 9 | prostate gland | 16 | 2 | 16 |
| 10 | testis | 35 | 5 | 19 |
| 11 | thyroid gland | 26 | 1 | 29 |

\*This data was obtained from the EMBL-EBI expression atlas (http://www.ebi.ac.uk/gxa).
Table that of the ovary which has highest TPM (82) and TPM cut-off is at least 1 for *SREBF1* and *RUNX3*. TPM: Transcript per million.

### Cell culture, transient transfections and reporter assays

Authenticated human HEK-293, HuH-7, Hep G2, rat BRL 3A and CHO cell lines were obtained from Indian national repository for cell lines and hybridomas at the National Centre for Cell Sciences, Pune, India and were cultured in Dulbecco’s Modified Eagle’s Medium (DMEM) with high glucose and glutamine (HyClone, Pittsburgh, USA), supplemented with 10% fetal bovine serum (Invitrogen, Waltham, USA), penicillin G (100 U/ml) and streptomycin sulfate (100 mg/ml) (Invitrogen, Waltham, USA) at 37°C with 5% CO_2_. These cell lines were routinely tested for mycoplasma infection and treated with BM-Cyclin (Merck, Burlington, USA) to eliminate mycoplasma, if detected. HEK-293, BRL 3A, HuH-7 and CHO cells at 60% confluence were transfected with 0.5-1.0 μg/well of either SHR-or WKY-*Hmgcr* promoter-reporter constructs or pGL3-Basic vector (as a control) and pCMV-βGal (β-galactosidase expression plasmid) for normalization. HEK-293, CHO and HuH-7 cells were transfected by calcium phosphate method (23) while BRL-3A cells were transfected by Targetfect F2 transfection reagent (Targeting Systems, El Cajon, USA). High quality endotoxin-free plasmids (prepared by HiPurA™ Endotoxin free Plasmid DNA Midiprep Purification Spin Kit; HiMedia, Mumbai, India) were used for all the transfections. In another set of experiments, 0.5-1.0 µg/well of SHR/WKY promoter mutants were also transfected in BRL 3A cells. pCMV-βGal (β-galactosidase expression plasmid) was used as internal control for transfection efficiency. After 24 hrs of transfection, cells were lysed for luciferase and β-gal assays. The luciferase assay was carried as previously described (24-26). The normalized promoter activities were expressed as luciferase/β-gal reading.

SHR/WKY-*Hmgcr* promoter reporter construct was co-transfected with different doses of pcDNA3.1-Flag-RUNX3 or SREBP1a (27) expression plasmid respectively. The total amount of plasmid DNA was made equal by an empty vector plasmid pcDNA3/CMV-7 control vector. Media was changed with fresh complete media in all the wells 7-8 hrs post-transfection. After 24-30 hrs of transfection, the cells were lysed for luciferase and Bradford (Bio-Rad, Hercules, USA) assays. The luciferase activities were expressed as luciferase activity/µg of protein.

For cholesterol depletion, BRL 3A cells (transfected with SHR/WKY-Hmgcr promoter construct) were treated with 5 mM of cholesterol-depleting reagent methyl-β-cyclodextrin (HiMedia, Mumbai, India) for 15 min. Cholesterol depletion was carried out in serum-free DMEM medium. Following cholesterol depletion, media was changed with fresh serum-free media and the cells were incubated for 6-9 hrs at 37°C in CO_2_ incubator. In another series of experiments, promoter construct (SHR/WKY-*Hmgcr*)-transfected BRL 3A cells were treated with exogenous cholesterol (20 µg/ml) in serum free media for 6-9 hrs. Following treatments, the cells were lysed for luciferase and Bradford assays.

### Isolation of RNA from SHR/WKY liver tissues and real-time PCR analysis

Total RNA was isolated from liver tissues of SHR and WKY (4-6 weeks old) using the TRIzol reagent (Invitrogen, Waltham, USA) followed by purification using Nucleospin columns (Machery-Nagel, Duren, Germany). cDNA synthesis was performed using High Capacity cDNA Reverse Transcription Kit (Applied Biosystems, Waltham, USA) and random hexamer primers. Quantitative real-time PCR (qPCR) was carried out using *Hmgcr* gene specific primers: forward, 5’-CCAATGTATCCGTTGTGGATCTG-3’ and reverse, 5’-GAGTTGCTGTTAAGTCACAG-3’ in an Eppendorf Mastercycler ep realplex real-time PCR system with DyNAmo ColorFlash SYBR Green qPCR Kit (Thermo Fisher, Waltham, USA). For normalization of *Hmgcr* expression, GAPDH abundance was measured using the following primers: forward, 5’-GACATGCCGCCTGGAGAAAC-3’ and reverse, 5’-AGCCCAGGATGCCCTTTAGT-3’. *Hmgcr* expression was normalized to GAPDH abundance and fold difference between SHR and WKY tissues was calculated by 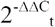 method (28).

### Chromatin Immunoprecipitation (ChIP) Assays

In order to confirm differential interactions of Runx3 and Srebf1 with the SHR-and WKY-*Hmgcr* promoters *in vivo* in context of chromatin, ChIP assays were carried out as described recently (29). In brief, BRL 3A cells were transfected with 10 µg of SHR/WKY-*Hmgcr* promoter constructs using Targetfect F2 transfection reagent (Targeting Systems, El Cajon, USA). Cells were crosslinked using 1% formaldehyde in PBS at room temperature for 12 min after 24 hrs of transfection. Chromatin was isolated and fragmented by sonication at 30% amplitude and 30 sec on-off pulse cycle. For immunoprecipitation reactions, precleared chromatin was incubated with 5 μg of ChIP grade antibodies i.e. anti-Runx3 (rabbit polyclonal, Santa Cruz Biotechnologies, Dallas, USA, #sc-30197X) and anti-SREBP1a (mouse monoclonal, Santa Cruz Biotechnologies, Dallas, USA, #sc-13551X) and pre-immune anti-rabbit IgG (Sigma-Aldrich, St. Louis, USA, I5006)/pre-immune anti-mouse IgG (Sigma-Aldrich, St. Louis, USA, I5831) over-night at 4°C. Immunoprecipitated complexes were reverse-crosslinked, digested with RNase and proteinase K followed by purification by HiPurA™ PCR product purification kit (HiMedia, Mumbai, India) and stored at −20°C until use. qPCR was carried out using immunoprecipitated chromatins to amplify DNA region encompassing the A-405G and -393_-382insTGCGGTCCTCC variations in the rat *Hmgcr* promoter using specific primers (FP: 5’-ACTACTTCCCCGCAGATCTC-3’ and RP: 5’-CCCCGAACTCAGAATTTGACC-3’) that amplify a 157 bp *Hmgcr* promoter domain. qPCR was performed by using QuantStudio™ 7 Flex Real-Time PCR System (Applied Biosystems, Waltham, USA) and DyNAmo ColorFlash SYBR Green qPCR Kit (Thermo Fisher, Waltham, USA) and results were expressed as fold enrichment over IgG signal or background.

### Western Blotting

For Western blot analysis, cells were lysed in radio-immunoprecipitation assay (RIPA) buffer [50 mM Tris–HCl (pH 7.2), 150 mM NaCl, 1% (v/v) Triton X-100, 1% (w/v) sodium deoxycholate, 1 mM EDTA and 0.1% (w/v) SDS] with 1 mM PMSF and protease inhibitor cocktail (Sigma-Aldrich, St. Louis, USA) followed by sonication. For tissue protein isolation from SHR/WKY liver tissues, 20-50 mg of liver tissue was homogenized in 0.5 ml RIPA buffer using a micropestle (Tarsons, Kolkata, India) in a 1.5 ml microcentrifuge tube. Following homogenization, the samples were sonicated, centrifuged at 14,000 rpm and the supernatant was stored in aliquots at −80°C. Bradford Assay (Bio-Rad, Hercules, USA) was used to estimate the protein concentrations in the cell or tissue lysates. Equal amount of protein samples (∼30-50 μg) were resolved by SDS-PAGE and transferred to either an activated PVDF (polyvinylidene difluoride; Pall Life Sciences, New York, USA) or a nitrocellulose membrane (Pall Life Sciences, New York, USA). The membranes were incubated with specific primary antibodies [anti-Hmgcr (abcam, Cambridge, USA, #ab174830) at 1:1000 dilution, anti-Runx3 (abcam, Cambridge, USA, #ab49117) at 1:1000 dilution, anti-Srebf1 (Santa Cruz Biotechnologies, Dallas, USA, #sc-13551) at 1:2500 dilution, anti-β-Actin (Sigma-Aldrich, St. Louis, USA, # A5441) at 1:7500 dilution and anti-Vinculin (Sigma-Aldrich, St. Louis, USA, # V9131) at 1:10000 dilution] overnight at 4°C following blocking with 3-5% of BSA/non-fat milk for 1 hr at room temperature. After 3 washes with 1xTBST, the membranes were incubated with HRP-conjugated secondary antibody specific for either rabbit (BioRad # 170-6515 at 1:3000 dilution for Hmgcr and Runx3) or mouse (Jackson Immunoresearch, West Groove, USA # 115-035-003 at 1:5000 dilutions for β-Actin, Vinculin and at 1:3000 for Srebf1) for 1 hr. The signals were detected using Clarity™ Western ECL Substrate kit (Bio-Rad, Hercules, USA) and a ChemiDoc Chemiluminescence Detection system (Bio-Rad, Hercules, USA). The intensities of signals/bands were quantified using ImageJ (30).

### Filipin staining of BRL 3A cells

BRL 3A cells (transfected with SHR/WKY-Hmgcr promoter construct) were treated with 20 µg/ml of exogenous cholesterol (HiMedia, Mumbai, India) or 5mM of cholesterol-depleting reagent methyl-β-cyclodextrin (MCD; HiMedia, Mumbai, India) for 6 hrs or 15 min, respectively. The media was changed to serum-free media following MCD treatment and the cells were incubated at 37 °C for 6-9 hrs. The cells were then washed with PBS and fixed with 3.6% formaldehyde in PBS for 10 min at room temperature. Following fixation, the cells were washed with PBS thrice and stained with 50 µg/ml of Filipin III (a fluorescent dye binding to free cholesterol) in the dark at room temperature for 3 hrs. The stained cells were imaged using an Olympus IX73 fluorescence microscope (Olympus, Tokyo, Japan).

### Analysis of serum cholesterol and systolic blood pressure values from various genetically-inbred hypertensive strains

The serum total cholesterol and systolic blood pressure data from different genetically-inbred hypertensive strains along with their respective normotensive control values was obtained from the National BioResource Project-Rat (http://www.anim.med.kyoto-u.ac.jp/nbr/) and PhenoMiner tool of the Rat Genome Database (http://rgd.mcw.edu/rgdweb/search/qtls.html?100). The identity of the strains (n=21) and related information is mentioned in Table 3.

**Table 3.**
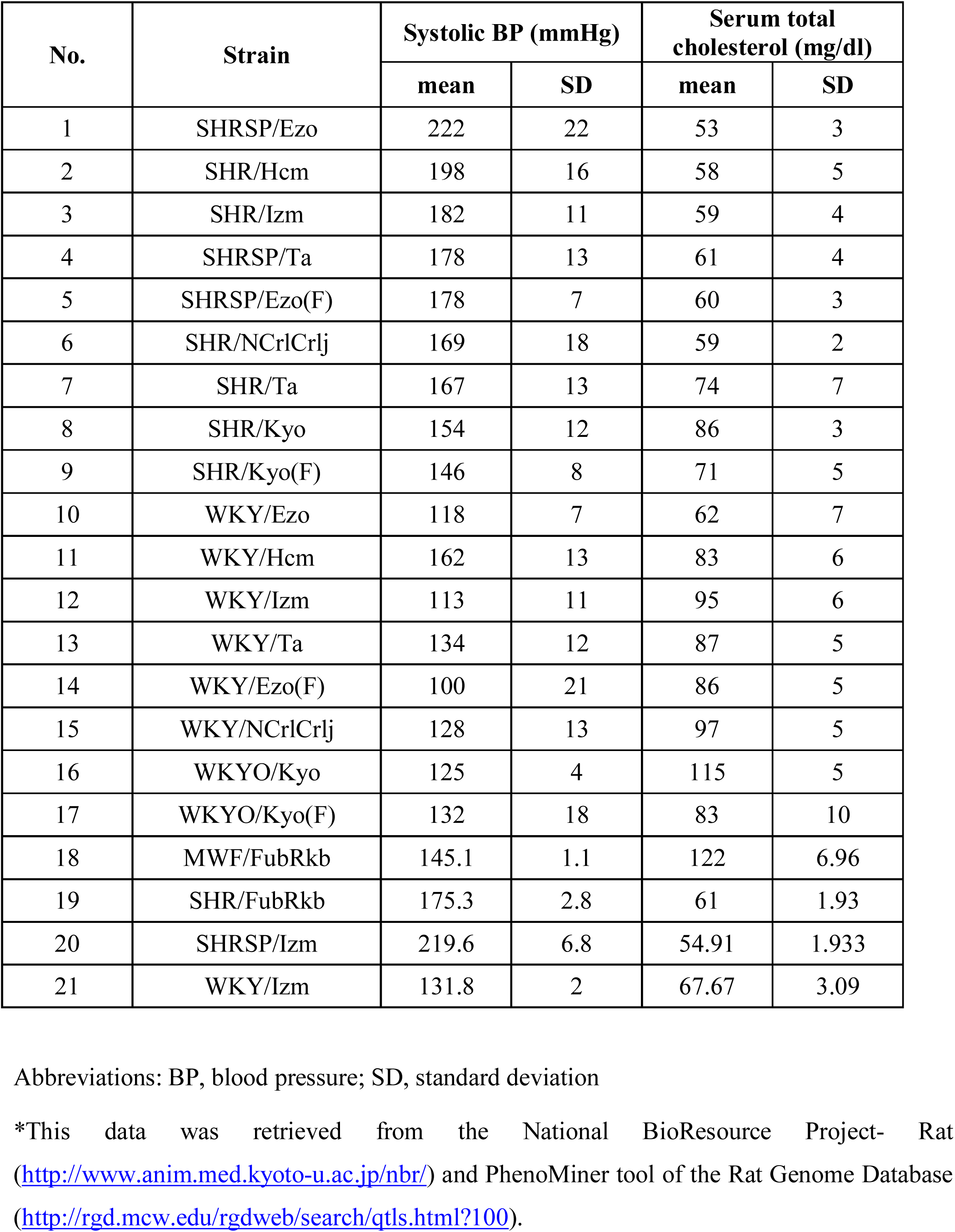
Systolic blood pressure and serum total cholesterol data from various genetically-inbred hypertensive rat strains with their respective controls*.

### Data presentation and statistics

All transient transfection experiments were performed at least three times and results were expressed as mean ± SEM of triplicates. Statistical significance was determined by Student’s t-test or one-way ANOVA with Newman-Keuls’s post-test, as appropriate for different experiments using Prism 5 program (GraphPad Software, San Diego, CA, USA). GraphPad Prism 5 software was used to generate all the graphs.

## RESULTS

### Comparative genomics analysis of human, mouse and rat *Hmgcr* gene sequences

Although various resources are available for genetic mapping in mice and mouse models are highly useful to dissect the genetic basis of complex human diseases (31), most of the quantitative trait loci (QTL) studies have been performed in rats because measurement of cardiovascular phenotypes in mice is technically challenging. We probed the Rat Genome Database (RGD) (15) for blood pressure QTLs on *Hmgcr*-harbouring chromosome 2 and detected several blood pressure QTLs (Fig.1A). The blood pressure-QTLs, their positions and respective LOD [logarithm (base 10) of odds] scores retrieved from RGD are shown in Table S1. Interestingly, the QTL Bp240 (45317613-103494306 bp) that harbours *Hmgcr* gene (27480226-27500654 bp, RGD ID: 2803) displayed the highest linkage (LOD score=4.0) with mean arterial blood pressure and systolic blood pressure (Fig.1A). In addition, QTLs-Bp243 (45317613-155139446 bp), Bp205 (45399390-62179292 bp) also housing the *Hmgcr* locus displayed significant linkage with diastolic blood pressure (LOD score=3.9) and mean arterial blood pressure (LOD score=3.476), respectively. Moreover, alignment of the mouse and human genomic regions with the orthologous rat sequences at *Hmgcr* locus using mVISTA showed >75% homology between the rodent sequences at exons, introns and untranslated regions (Fig.1B); the human genomic regions show high conservation at the 5’ UTR and exonic regions with the rat sequences. In general, the extent of homology between each of the twenty *Hmgcr* exons in mouse and rat was higher (>85%) than the noncoding regions (Fig.1B). Thus, rat *Hmgcr* appeared as a logical candidate gene for studying the mechanisms of hypertension.

**Fig.1.**
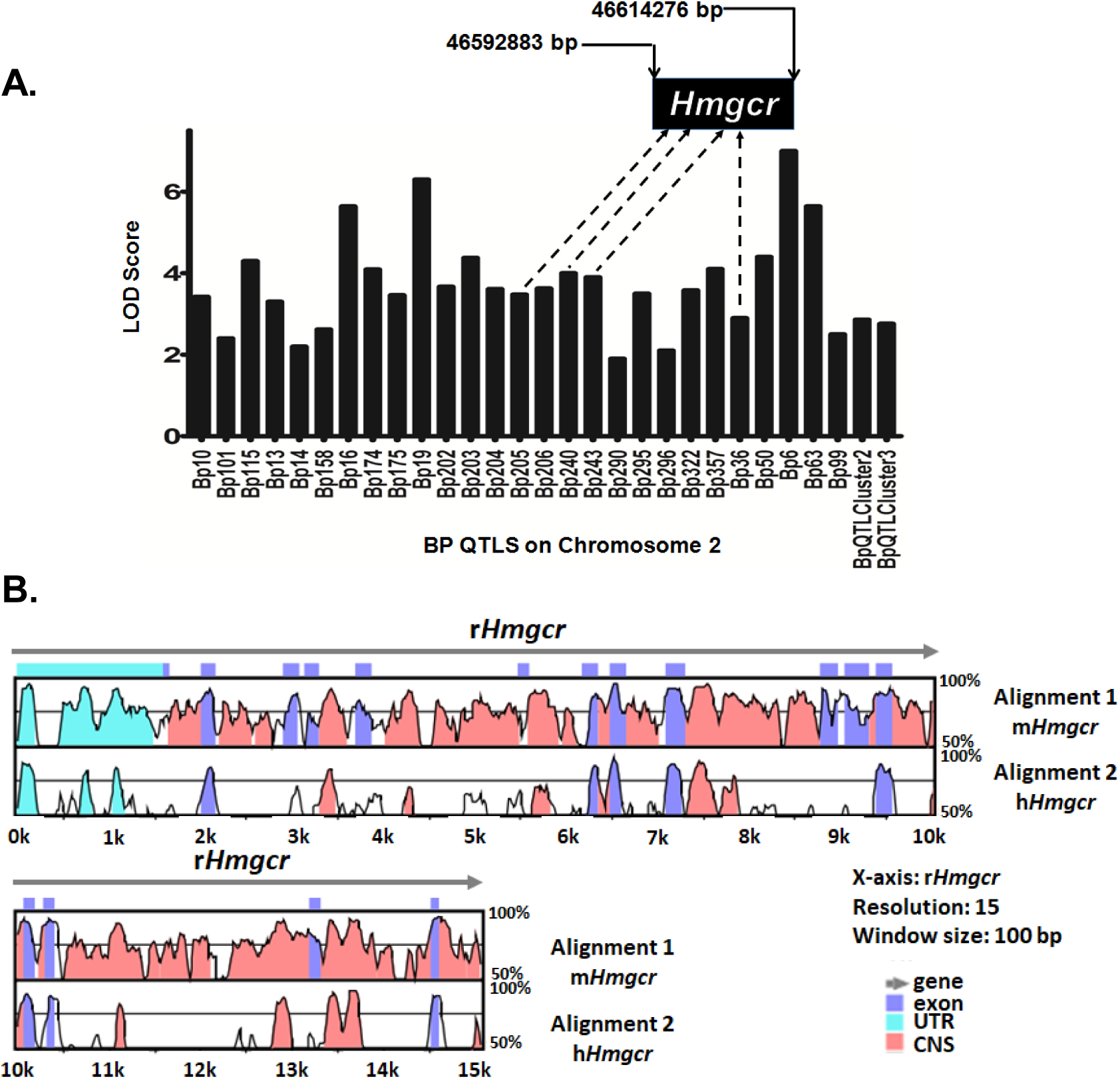
Graphical representation of rat blood pressure-QTLs and homology between mouse-, human-and rat-*Hmgcr* gene sequences. (A)The blood pressure-QTLs and their respective LOD scores (retrieved from Rat Genome Database) were plotted. The genomic position of rat *Hmgcr* (*rHmgcr*) gene is indicated. (B) Conservation analysis of human/mouse and rat *Hmgcr* sequences was performed using mVISTA. The horizontal axis represents the *rHmgcr* gene (chr2:27480226-2750064) as the reference sequence, whereas the upper vertical axis indicates the percentage homology between rat and mouse *Hmgcr* gene (chr13:96,650,579-96,666,685); the lower vertical axis indicates the percentage homology between rat and human *Hmgcr* (*HMGCR*) gene (chr5:75,337,168-75,362,104). The length of comparison or window size was set to 100 bp with a minimum of 70% match. The annotation of gene is represented by different colors. Mouse/rat/human *Hmgcr* genes comprise of twenty exons; 5’-UTR is not visible in the homology plot due to their very small sizes. CNS: conserved non-coding sequences; UTR: Untranslated region.

### *Hmgcr* is differentially expressed in SHR and WKY liver tissues

To investigate whether the expression of Hmgcr differs between SHR and WKY in liver tissues, we assessed the Hmgcr mRNA and protein levels in the liver (primary site for cholesterol biosynthesis). Our qPCR analysis revealed that the Hmgcr transcript levels were elevated in liver tissues of WKY (∼4-fold, p<0.05; Fig.2A) as compared to SHR. In corroboration, the Hmgcr protein level were also higher in WKY liver issues as compared to SHR (∼by 2-fold, p<0.05; Fig.2B).

**Fig.2.**
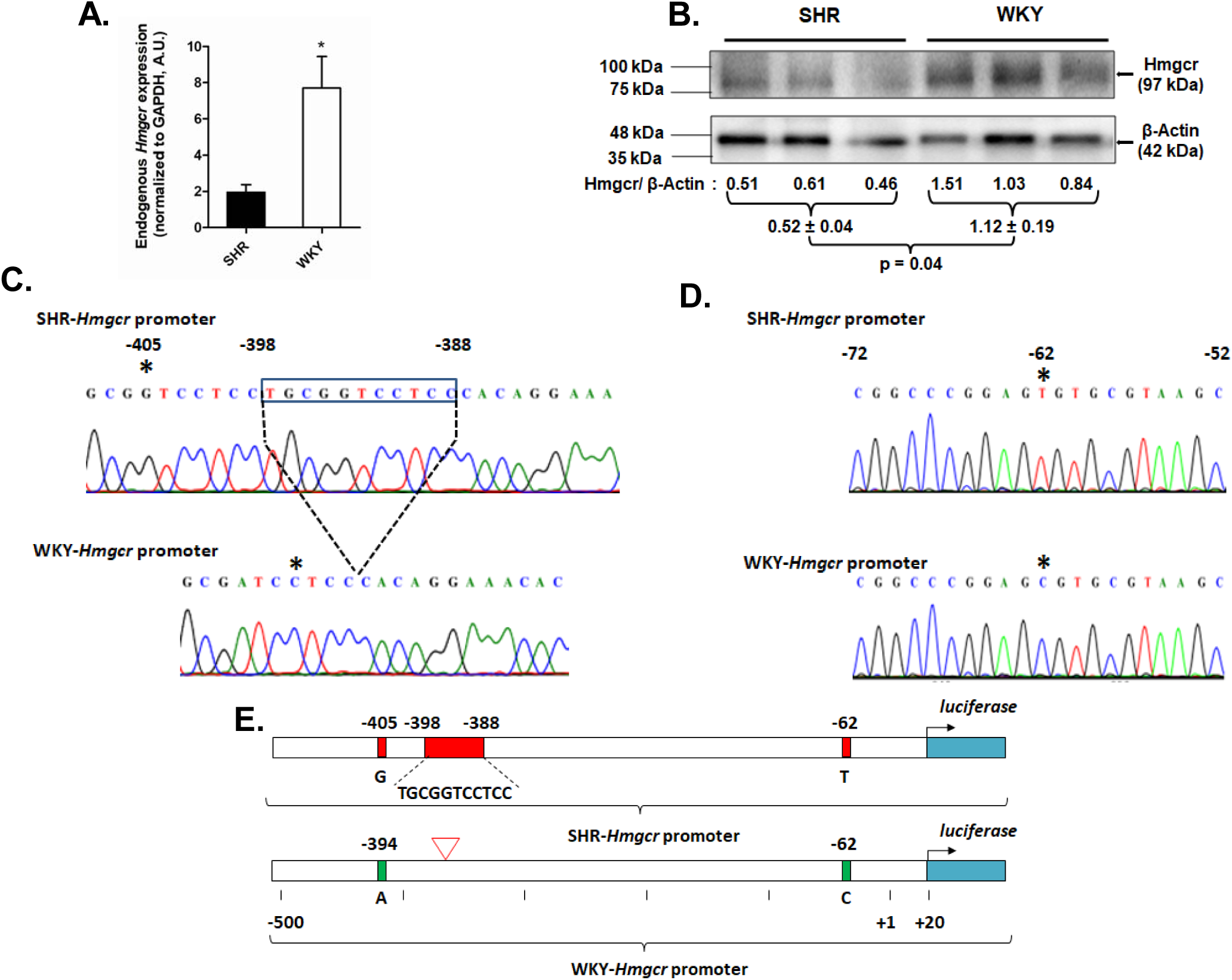
Endogenous hepatic expression of Hmgcr and *Hmgcr* promoter polymorphisms in SHR and WKY. Total RNA and protein were isolated from liver tissues (n=4) of SHR and WKY rats. RNA was reverse transcribed and the cDNA was used as a template for (A) qPCR using gene specific primers for *Hmgcr* and GAPDH. *Hmgcr* expression was normalized to GAPDH. (B) Representative Western blot analysis for Hmgcr in liver tissues of SHR and WKY. The relative levels of Hmgcr are indicated below the Western blot images. The SHR/WKY-*Hmgcr* promoter regions were sequenced as described in the materials and methods section. Representative chromatograms showing (C) A-405G and -393_-382insTGCGGTCCTCC and (D) C-62T variations in SHR *Hmgcr*-promoter with respect to WKY-*Hmgcr* promoter domains. The nucleotides are numbered with respect to the first nucleotide of cap site as +1 in WKY-*Hmgcr* promoter. Star indicates nucleotide change while the box indicates insertions. (E) Schematic representation of *Hmgcr* (SHR and WKY) promoter reporter constructs harbouring the identified variations. Statistical significance was determined by Student’s *t*-test (unpaired, 2-tailed). *p<0.05 as compared to SHR.

### Identification of variations in the SHR/WKY-*Hmgcr* promoter domain and conservation of nucleotide sequences at the polymorphic regions among different mammals

To unravel the molecular mechanism of differential expression of *Hmgcr* between SHR and WKY, the proximal ∼1 kb region of the *Hmgcr* promoter domain was sequenced in these models. ClustalW (17) alignment of the SHR-and WKY-*Hmgcr* promoter sequences revealed three variations: A-405G, C-62T and a 11 bp insertion (-393_-382insTGCGGTCCTCC) in SHR-*Hmgcr* promoter with respect to WKY-*Hmgcr* promoter (Fig.2E).

To analyse if there is any conservation across the polymorphic domains of the rat *Hmgcr* promoter with orthologous *Hmgcr* sequences in various mammalian species, the SHR/WKY promoter sequences were aligned with the *Hmgcr* promoter sequences of various mammals (mouse, human, rat, orangutan, pig and cow) using the GeneDoc software (22). Interestingly, the G-allele at −405 bp position discovered in SHR was conserved at the corresponding positions in *Hmgcr* promoter of mouse, human, orangutan, pig and cow (Fig.3A). However, the -393_-382insTGCGGTCCTCC was observed only in SHR among the sequences included (Fig.3A). Additionally, the T-allele at the −62 bp position in SHR was absent in the other mammals included in this analysis (Fig.3B).

**Fig.3.**
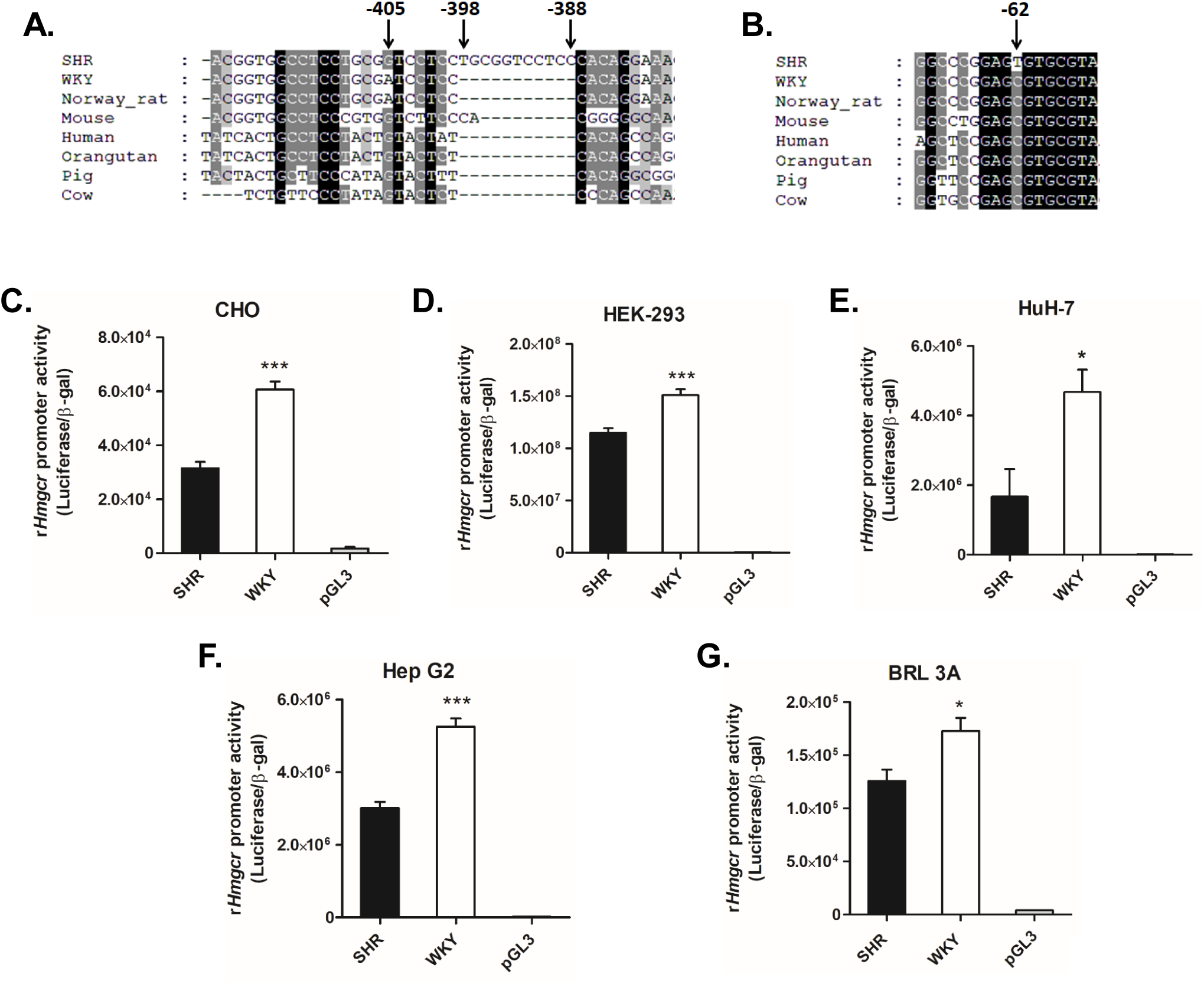
Differential expression of *Hmgcr*-promoter reporter constructs in cultured cells. Conservation of the *Hmgcr* promoter domains harboring the (A) A-405G, -393_-382insTGCGGTCCTCC and (B) C-62T polymorphisms among various mammals. The location of the variations between SHR-*Hmgcr* and WKY-*Hmgcr* are indicated by arrows with numbers. (C-H) Comparison of *Hmgcr*-promoter strengths (SHR and WKY) in various cultured cells. The *Hmgcr*-promoter reporter constructs (SHR/WKY) or pGL3-Basic vector were transiently transfected with co-transfected control plasmid pCMV-βGal into (C) CHO, (D) HEK-293, (E) HuH-7, (F) Hep G2 and (G) BRL 3A cells. The transfected cells were assayed for luciferase and β-galactosidase activity 24-30 hrs post-transfection. The promoter activities were normalized to the β-galactosidase and expressed as mean ± SEM of triplicate values. Statistical significance was determined by one-way ANOVA with Newman-Keuls multiple comparison test. *p<0.05 and ***p<0.001 as compared to the SHR-*Hmgcr* promoter activity.

### Differential activities of *Hmgcr* promoters under basal conditions

To investigate the basal promoter activity of SHR/WKY-*Hmgcr* domains, the SHR and WKY-*Hmgcr* promoter constructs as well as pGL3-Basic plasmid (as a control) were transfected into CHO, HEK-293, HuH-7, Hep G2 and BRL 3A cells (Fig. 3C-G). In all the cell lines tested, the WKY-*Hmgcr* promoter activity was significantly higher as compared to the SHR-*Hmgcr* promoter: ∼2-fold in CHO (p<0.0001), ∼1.3-fold in HEK-293 (p<0.0001), ∼2.8-fold in HuH-7 (p<0.05), ∼1.75-fold in Hep G2 (p<0.0001) and ∼1.4-fold in BRL 3A (p<0.05). Thus, consistent with the Hmgcr mRNA and protein expression in SHR/WKY liver tissues, the WKY-*Hmgcr* promoter showed enhanced promoter activity as compared to the SHR-*Hmgcr* promoter.

### Functional consequences of Hmgcr promoter variations and putative transcription factors regulating *Hmgcr* expression

To access if the variations identified in the SHR/WKY-Hmgcr domains were functional, various *Hmgcr* promoter mutants were generated by site-directed mutagenesis (Fig.4A). The *Hmgcr* promoter constructs (viz. SHR, WKY, SDM, W394, S62, W62 and S405) and the pGL3-basic plasmid (as control) were transfected into BRL 3A and HuH-7 cells (Fig.4). Interestingly, the W394 and S62 constructs showed diminished promoter activity as compared to the WKY-*Hmgcr* promoter in both BRL 3A and HuH-7 cell lines (p<0.01). This suggests that the G allele at −405 bp position and -393_-382insTGCGGTCCTCC in SHR-*Hmgcr* promoter may contribute to the diminished promoter activity of SHR-*Hmgcr* promoter. However, the contribution of the variation at the −62 bp to the overall basal *Hmgcr* promoter activity of SHR or WKY was uncertain as both S62 and W62 constructs showed no significant difference with SHR and WKY-*Hmgcr* promoter activity, respectively, in both the cell lines tested. Therefore, differential activities of SHR/WKY-*Hmgcr* promoters may be due to the allele at −405 bp position and the insertion at -393/-382 bp position.

**Fig.4.**
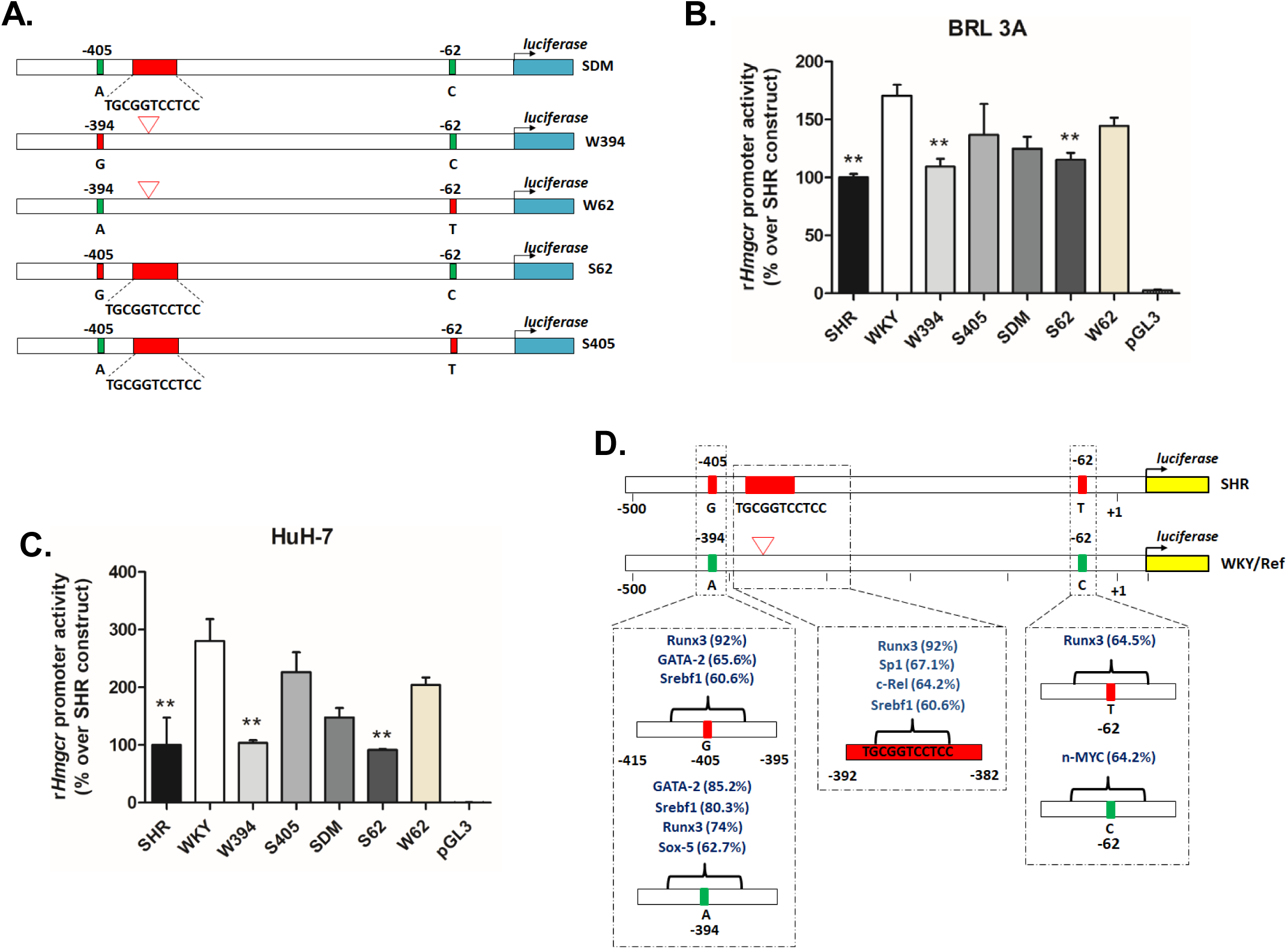
Functional implications of *Hmgcr*-promoter variations in cultured hepatocytes. (A) Schematic representations of *Hmgcr*-promoter mutant constructs generated by site-directed mutagenesis using SHR/WKY-*Hmgcr* promoter construct as template. The *Hmgcr*-promoter reporter constructs (SHR/WKY), promoter mutants and pGL3-Basic were co-transfected with pCMV-βGal plasmid into (B) BRL 3A, (C) HuH-7 cells. Promoter activities were normalized to the β-galactosidase and expressed as % over SHR-*Hmgcr* promoter activity. The results are mean ± SEM of triplicate values. Statistical significance was determined by one-way ANOVA with Newman-Keuls multiple comparison test. **p<0.01 as compared to the WKY-*Hmgcr* promoter activity. (D) Schematic representation of putative transcription factor binding sites across the *Hmgcr* promoter variations in SHR and WKY-*Hmgcr* promoter constructs. Multiple *in silico* tools were used to predict putative transcription factor sites and detailed in the materials and methods section. Only transcription factor sites predicted by at least two programs are shown and the TFSEARCH scores are indicated in the brackets.

Since we identified variations in the SHR/WKY-*Hmgcr* promoter domains, we carried out stringent *in silico* analysis to identify potential transcription factors that may alter the expression of Hmgcr in these models of EH. Computational analyses were carried out using four different tools to predict the putative transcription factor binding sites: ConSite (18), TFSEARCH (19), P-Match (20) and LASAGNA (21). Putative transcription factors predicted by at least two programs for different scores of binding across the discovered variations were shortlisted and shown in Fig.4D. Additionally, only those transcription factors predicted to bind to the functional variations (viz. A-405G and -393_-382insTGCGGTCCTCC) were chosen for further validation. Interestingly, these predictions revealed preferential binding of Runx3 at all three discovered variations in SHR as compared to WKY-*Hmgcr* promoter while Srebf1 showed higher binding scores to the A allele at -394 bp position in the WKY-*Hmgcr* promoter and the insertion at -393/-382 bp in SHR-*Hmgcr*. Hence, Runx3 and Srebf1 were selected for further experimental studies.

### Important roles of Runx3 and Srebf1 in transcriptional regulation of Hmgcr

To test the effect of predicted transcription factors (Runx3 and Srebf1) on *Hmgcr* gene regulation, we carried out co-transfection experiments in BRL 3A cells. Over-expression of Runx3 caused a dose-dependent reduction in the SHR-*Hmgcr* promoter activity (∼up to 2-fold, p<0.01); the WKY-*Hmgcr* activity was also dose-dependently diminished (∼up to 1.8∼fold, p<0.001) under similar conditions (Fig.5A). The SHR-*Hmgcr* promoter activity, in general, was lower than the WKY-*Hmgcr* promoter activity at each concentration of Runx3 (Fig.5A). To access the extent of interactions of Runx3 with the SHR and WKY-*Hmgcr* promoters *in vivo* in the context of chromatin, ChIP assays were performed in HuH-7 cells transfected with SHR-and WKY-*Hmgcr* constructs. qPCR for chromatin immunoprecipitated with Runx3 antibody revealed significant enrichment of SHR-*Hmgcr* domain (by ∼2.2-fold, p<0.0001 with respect to IgG) and WKY-*Hmgcr* domain (by ∼1.4-fold, p<0.01 with respect to IgG) (Fig.5B). Moreover, the fold-enrichment of the SHR-*Hmgcr* domain was significantly higher for immunoprecipitation with Runx3 as compared to WKY-*Hmgcr* domain (by ∼1.5-fold, p<0.01; Fig.5B).

**Fig.5.**
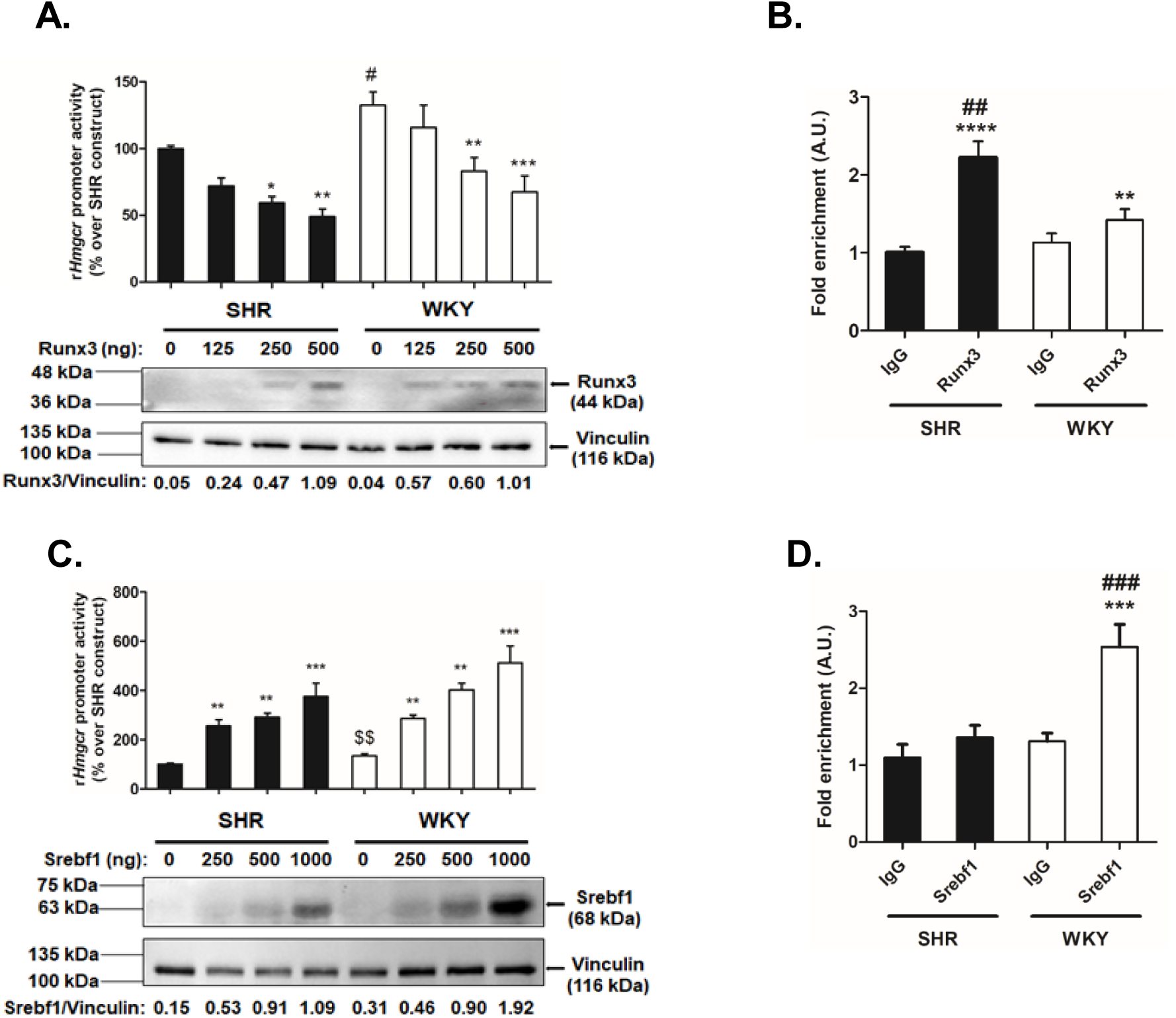
Role of Srebf1 and Runx3 in transcriptional regulation of Hmgcr. (A) Modulation of SHR/WKY-*Hmgcr* promoter activity by ectopic expression of Runx3. BRL 3A cells were co-transfected with 500 ng/well of *Hmgcr*-promoter constructs (SHR/WKY) and increasing doses of pcDNA3.1-Flag-RUNX3 plasmid. The luciferase activity was normalized to total protein and expressed as % over SHR-*Hmgcr* promoter activity. The results are mean ± SEM of triplicate values. Statistical significance was determined by one-way ANOVA with Newman-Keuls multiple comparison test. *p<0.05, **p<0.01 and ***p<0.001 as compared to the basal condition (without a co-transfected pcDNA3.1-Flag-RUNX3 plasmid), ^#^p<0.05 as compared to the basal SHR-*Hmgcr* promoter activity. (B) ChIP assay for endogenous 0interaction of Runx3 with *Hmgcr* promoter domains. Chromatin isolated from HuH-7 cells (transfected with SHR/WKY-*Hmgcr* promoter constructs) was immunoprecipitated by anti-Runx3 antibody or pre-immune IgG. qPCR was carried out with DNA purified from respective cocktails and primers flanking the A-405G and -393_-382insTGCGGTCCTCC variations. The results are expressed as fold enrichment over pre-immune IgG. Statistical significance was determined by one-way ANOVA with Newman-Keuls multiple comparison test. **p<0.01 and ****p<0.0001 as compared to the corresponding IgG condition, ^##^p<0.01 as compared to corresponding WKY-condition. (C) Effect of over-expression of Srebf1 on *Hmgcr*-promoter activity. BRL 3A cells were transiently transfected with increasing concentration of SREBP1a expression plasmid and 500 ng/well of SHR/WKY-*Hmgcr* promoter constructs. The normalized luciferase activity was expressed as % over SHR-*Hmgcr* promoter activity and results. are mean ± SEM of triplicate values. Statistical significance was determined by one-way ANOVA with Newman-Keuls multiple comparison test and Student’s *t*-test (unpaired, 2-tailed). **p<0.01 and ***p<0.001 as compared to the basal condition (without a co-transfected SREBP1a expression plasmid), ^$$^p<0.01 as compared to the basal SHR-*Hmgcr* promoter activity (Student’s *t*-test). (D) *In vivo* interactions of Srebf1 with SHR/WKY-*Hmgcr* promoter domain in the context of chromatin. ChIP assay of HuH-7 cells transfected with SHR/WKY-*Hmgcr* promoter constructs was performed using anti-Srebf1 antibody/pre-immune IgG. The purified, immunoprecipitated chromatins were subjected to qPCR using primers flanking the A-405G and -393_-382insTGCGGTCCTCC variations. Fold-enrichment over IgG is shown. Statistical significance was determined by one-way ANOVA with Newman-Keuls multiple comparison test. ***p<0.001 as compared to the corresponding IgG condition, ^###^p<0.01 as compared to corresponding SHR-condition.

Similarly, co-transfection of Srebf1 augmented the SHR-*Hmgcr* promoter activity (up to ∼3.7-fold, p<0.001) and WKY-*Hmgcr* promoter activity (up to ∼3.8-fold, p<0.001) in a concentration-dependent manner (Fig.5C). Interestingly, ChIP assays displayed enhanced enrichment of WKY-*Hmgcr* domain (up to ∼2-fold, p<0.001 with respect to IgG) for immunoprecipitation with Srebf1 (Fig.5D). In contrast, there was no significant enrichment of the SHR-*Hmgcr* domain with respect to IgG (Fig.5D). Taken together, these results suggest that Runx3 exerts stronger interactions with SHR-*Hmgcr* promoter than the WKY-*Hmgcr* promoter while Srebf1 binds to WKY-*Hmgcr* promoter with higher affinity as compared to the SHR-*Hmgcr* promoter.

### Association of Hmgcr expression with Runx3 and Srebf1 levels in rat models of hypertension and human tissues

In view of our findings of modulation of *Hmgcr* promoter activity by Runx3 and Srebf1, we investigated the Runx3 and Srebf1 protein levels in liver tissues of SHR and WKY. Western blot analysis showed enhanced Runx3 protein levels (up to ∼5-fold, p<0.0001) in SHR liver tissues as compared with WKY (Fig.6C). On the other hand, the Srebf1 protein levels were reduced (up to ∼8-fold, p<0.0001) in SHR as compared to WKY in the liver (Fig.6D).

**Fig.6.**
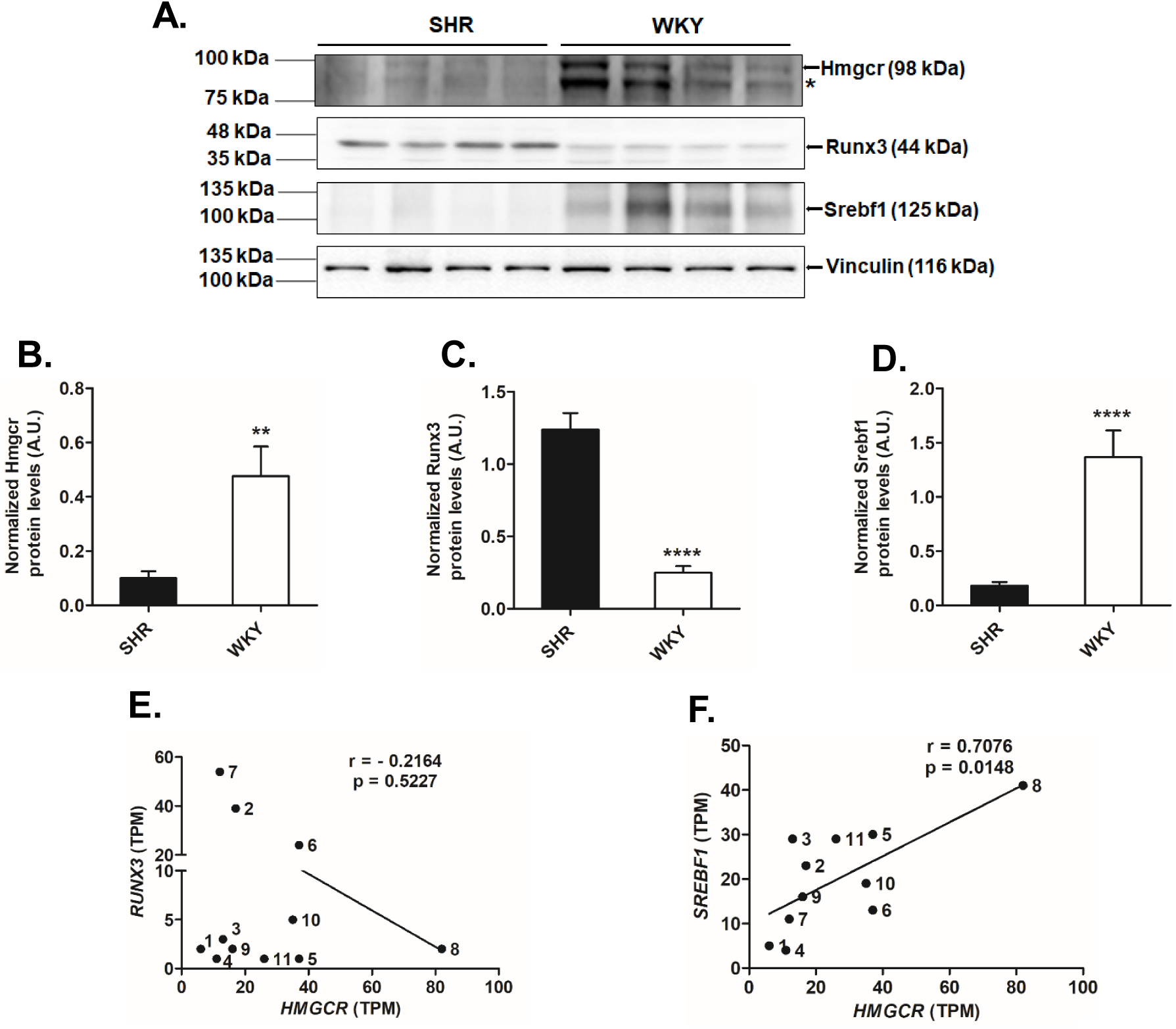
Endogenous expression of Hmgcr, Runx3 and Srebf1 in tissue samples of rat models of hypertension and humans. (A) Representative Western blot image depicting the endogenous Hmgcr, Runx3 and Srebf1 protein levels in liver tissues of SHR (n=4). *indicates a non-specific band. Quantification of endogenous (B) Hmgcr, (C) Runx3 and (D) Srebf1 protein levels in liver tissues of SHR/WKY (n=3). The Hmgcr/Runx3/Srebf1 protein levels were normalized to vinculin from three independent blots and shown as a bar plot. Statistical significance was determined by Student’s *t*-test (unpaired, 2-tailed). **p<0.01 and ****p<0.0001 as compared to SHR. (E) Correlation between transcript levels of (E) HMGCR and RUNX3 and (F) HMGCR and SREBF1 in various human tissues [n=12; the identity of samples (indicated by numbers) is provided in Table 2] using data retrieved from EMBL-EBI expression atlas (http://www.ebi.ac.uk/gxa).

In order to extent our findings to human tissues, correlation analysis was performed between HMGCR and RUNX3/Srebf1 transcript levels in various human tissues using two expression databases. The transcript level of HMGCR showed no correlation to the *RUNX3* expression in human tissues (Fig.6E, S1A). However, correlation analysis between transcript levels of HMGCR and SREBF1 across various human tissues revealed a significant positive correlation (Pearson *r* = 0.6395, p = 0.02 for EMBL-EBI expression data, Fig.6F; Pearson *r* = 0.5437, p = 0.003 for GTEx Portal data, Fig.S1B) suggesting a key role of Srebf1 in *HMGCR* regulation.

### Intracellular cholesterol modulates *Hmgcr* expression via Runx3 and Srebf1

As Hmgcr protein level is influenced by intracellular cholesterol level and regulated by a negative feed-back mechanism (6) we sought to determine whether modulating the endogenous cholesterol level affects SHR/WKY-*Hmgcr* promoter activity. Accordingly, BRL 3A cells transfected with SHR-and WKY-*Hmgcr* promoter constructs were treated with either cholesterol (20 µg/ml) or methyl-β-cyclodextrin (MCD, a cyclic oligosaccharide that reduces intracellular cholesterol level; 5 mM) (Fig.7A, B). Indeed, exogenous cholesterol treatment diminished both the SHR (by ∼1.9-fold, p<0.05) and WKY-*Hmgcr* promoter activities (by ∼3-fold, p<0.0001) (Fig.7A). In corroboration, intracellular cholesterol depletion enhanced both the SHR (by ∼2.5-fold, p<0.05) and WKY-*Hmgcr* promoter activities (by ∼1.62-fold, p<0.05) (Fig.7B). The increase/decrease in the intracellular cholesterol level upon cholesterol treatment/depletion experiments were confirmed by Filipin staining (Fig.S2). Western blot analysis revealed that the endogenous Hmgcr protein level decreased (up to ∼ 1.35-fold, p<0.05)/increased (up to ∼ 1.3-fold, p<0.01) upon cholesterol treatment/depletion, respectively (Fig.7C, D). Interestingly, Runx3 protein level was elevated (up to ∼ 1.6-fold, p<0.05)/reduced (up to ∼ 1.75-fold, p<0.05) upon cholesterol treatment/depletion. (Fig.7E). On the other hand, Srebf1 protein level was diminished (up to ∼ 1.4-fold, p<0.05)/augmented (up to ∼ 1.7-fold, p<0.05) upon cholesterol/MCD treatment, respectively (Fig.7F). These results suggest that intracellular cholesterol level regulates SHR/WKY-*Hmgcr* promoter expression via Runx3 and Srebf1.

**Fig.7.**
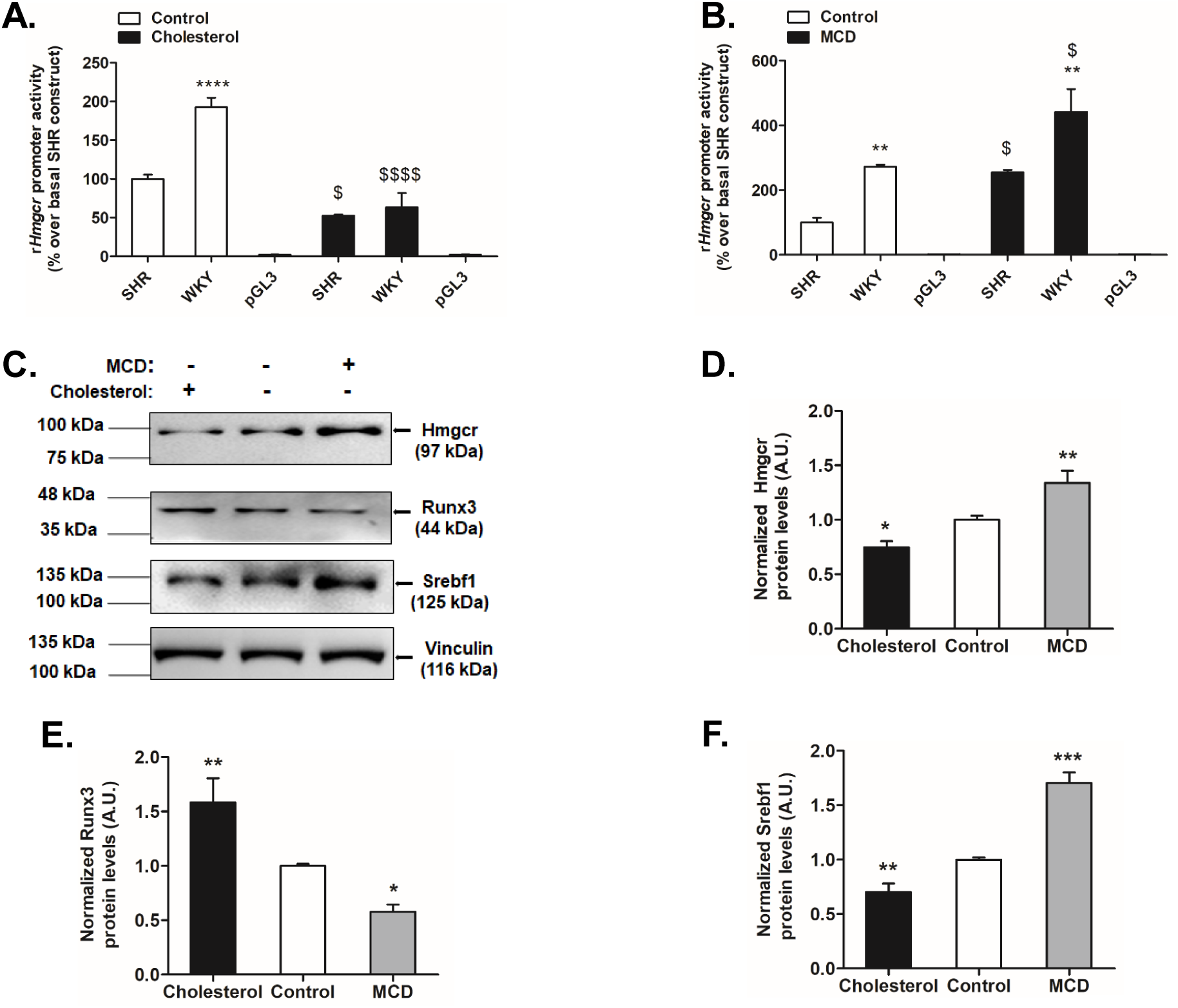
Intracellular cholesterol modulates expression of *Hmgcr* promoter via Runx3 and Srebf1. BRL 3A cells were transfected with SHR/WKY-*Hmgcr* promoter constructs and 24 hours post-transfection cells were treated with (A) either 20 µg/ml of cholesterol ((3β)-cholest-5-en-3-ol) or (B) 5 mM of cholesterol-depleting reagent methyl-β-cyclodextrin (MCD) for 6 hrs or 15 minutes, respectively. After incubation for 6-9 hrs in serum free media, the cells were assayed for luciferase and Bradford assay. The luciferase activity was normalized to total protein and expressed as % over basal SHR-*Hmgcr* promoter activity. Statistical significance was determined by one-way ANOVA with Newman-Keuls multiple comparison test. **p<0.01 and ****p<0.0001 as compared to the corresponding SHR-condition, ^$^p<0.05 and ^$$$$^p<0.0001 with respect to corresponding basal condition. (C) Western blot analysis of total protein isolated from BRL 3A cells treated with cholesterol or MCD was carried out to determine endogenous Hmgcr, Runx3 and Srebf1 levels. Quantification of endogenous (D) Hmgcr, (E) Runx3 and (F) Srebf1 protein levels in BRL 3A cells upon cholesterol/MCD treatment (n=3). The protein levels of Hmgcr, Runx3 and Srebf1 were normalized to vinculin from three independent blots and shown as a bar plot. Statistical significance was determined by one-way ANOVA with Newman-Keuls multiple comparison test. *p<0.05, **p<0.01 and ***p<0.001 as compared to the basal condition.

### Inverse correlation of serum cholesterol levels and systolic blood pressure in rat models of hypertension

Consistent with diminished *Hmgcr* promoter activity and Hmgcr mRNA/protein expression in SHR as compared to WKY, the serum cholesterol level of SHR is also reduced in comparison to WKY (Fig.S3). In order to test if other genetically-inbred hypertensive rat strains also show diminished serum cholesterol levels with respect to their controls, we retrieved serum total cholesterol and systolic blood pressure values of 21 strains from the National BioResource Project-Rat (http://www.anim.med.kyoto-u.ac.jp/nbr/) and PhenoMiner tool of the Rat Genome Database (http://rgd.mcw.edu/rgdweb/search/qtls.html?100) (Table 3). Interestingly, the serum cholesterol level and systolic blood pressure showed a significant negative correlation (Pearson *r* = −0.6475, p = 0.0015; Fig.S4) across various genetic rat models of hypertension suggesting that reduction in serum cholesterol levels might be a common compensatory mechanism in response to elevated blood pressure in these strains.

## DISCUSSION

### Overview

Dyslipidemia is a strong predictor of cardiovascular diseases including hypertension, obesity, atherosclerosis, coronary artery disease, and type 2 diabetes (32-34). HMGCR, a key regulator of dyslipidemia, is targeted by statins to lower high cholesterol levels and risk of CVDs (35, 36). Several studies also reported associations of single nucleotide polymorphisms (SNPs) in the *HMGCR* locus (viz. rs12654264, rs3846662, rs7703051, rs5908 rs12916, rs17238540 and rs3846663) with level of total cholesterol, LDL cholesterol and risk for dyslipidemia/stroke/blood pressure and coronary heart disease (37-44). Despite such associations of *Hmgcr* with lipid homeostasis and cardiovascular pathophysiology, the transcriptional regulation of *Hmgcr* in rat models of essential hypertension which are routinely used to investigate the molecular mechanisms underlying CVDs, has not been investigated. In this study, we found that Hmgcr is differentially expressed in liver tissues of these hypertensive rat models. Furthermore, sequencing and systematic experimental analyses revealed functional genetic variations (viz. A-405G and -393_-382insTGCGGTCCTCC) that altered the binding affinities of Runx3 and Srebf1 for SHR and WKY-*Hmgcr* promoters thereby resulting in diminished SHR-*Hmgcr* promoter activity as compared to WKY (Fig.3). The variant G allele at −405 bp in SHR was also detected in corresponding *Hmgcr* sequences of various mammals suggesting the preservation/occurrence of this variation in the course of evolution (Fig.3).

### Molecular basis of modulated Hmgcr expression: Role of Runx3 and Srebf1

In the view of differential expression of Hmgcr in SHR/WKY liver tissues (Fig.2), do variations A-405G and -393_-382insTGCGGTCCTCC alter the *Hmgcr* promoter activity? Computational analysis revealed that the G allele at the −405 bp and insertion at -393/382 bp in SHR-*Hmgcr* promoter offered better binding sites for Runx3 as compared to the A allele (Fig.4D). On the other hand, the A allele at -394 bp in WKY-*Hmgcr* promoter harboured a stronger Srebf1 binding site as compared to the G allele.

Runt-related transcription factor 3 (Runx3), a transcription factor belonging the Runt-related transcription factor family, is abundantly expressed in the hematopoietic system and involved in a variety of physiological processes including B-cell/T-cell differentiation, development of gastrointestinal tract and dorsal root ganglia neurogenesis (45, 46). Consistent with our *in silico* predictions, over-expression of Runx3 reduced the SHR-*Hmgcr* promoter activity to a greater extent than the WKY-*Hmgcr* promoter activity (Fig.5A). In corroboration, ChIP assays revealed Runx3 binds with higher affinity to the SHR-*Hmgcr* promoter than the WKY-*Hmgcr* promoter domain (Fig5B). In line with our finding of Runx3 as a repressor of SHR/WKY-*Hmgcr* promoter activity, various reports indicate the inhibitory role of Runx3 in transcriptional regulation (47-49). Moreover, Western blot analysis revealed elevated Runx3 protein level in SHR liver tissues than WKY (Fig.6C) suggesting that higher Runx3 level coupled with enhanced Runx3 binding to the SHR-*Hmgcr* promoter may contribute to the overall decrease in the Hmgcr expression in liver tissues of SHR as compared to WKY. Thus, this study, for the first time to our knowledge, revealed the role of Runx3 in the regulation of *Hmgcr* expression.

In line with the reported role of Srebf1 in *Hmgcr* expression (50) and our computational analyses, ectopic expression of Srebf1 caused a pronounced enhancement in the WKY-*Hmgcr* promoter activity than the SHR-*Hmgcr* promoter (Fig.5C). The differential binding of Srebf1 was confirmed by ChIP assays (Fig.5D). Sterol-regulatory element-binding protein 1 (SREBP-1) or Srebf1, encoded by *Srebf1* gene, has two protein isoforms that are well known transcription factors regulating hepatic cholesterol and fatty acid synthesis (50). Of note, over-expression of SREBPs in rodents causes hepatic steatosis and a transgenic SHR strain over-expressing human SREBP-1a exhibits multiple features of metabolic syndrome including hypertriglyceridemia, hyperinsulinemia and hypertension along with hepatic steatosis (51). Interestingly, genetic variations in SREBP have been reported to be associated with various cardiovascular complications in humans (52, 53). Does SHR show diminished hepatic level of Srebf1 contributing to the overall decrease in the Hmgcr protein and mRNA levels? Indeed, Western blot analysis reveals reduced Srebf1 level in SHR than WKY liver tissues (Fig.6A, D). This is in corroboration with a study that reported genetic variants in *Srebf1* gene in SHR account for lower Srebf1 mRNA/protein levels and *Srebf1* promoter activity (54).

In light of the role of Runx3 and Srebf1 in the transcriptional regulation of rat *Hmgcr*, we asked if the levels of Hmgcr and these transcription factors in human tissues correlated? Indeed, the HMGCR and SREBF1 transcript levels were significantly positively correlated across various tissues of human origin (Fig. 5F S3B) suggesting that SREBF1 could regulate HMGCR expression in human tissues. On the other hand, RUNX3 transcript levels showed a negative correlation with HMGCR transcripts levels which was not significant (Fig.6E). This observation is consistent with the dual role of RUNX3 as an activator as well as a repressor in different tissues (47-49, 55-57)

### Regulation of Hmgcr expression under pathophysiological conditions by Runx3 and Srebf1

Sterols regulate *HMGCR* expression via transcriptional, post-transcriptional and post-translational mechanisms (58). In brief, elevated sterols inhibit Sterol Regulatory Element Binding Proteins (SREBPs) thereby, diminishing *HMGCR* expression (59). The post-transcriptional and post-translational regulation mechanisms of HMGCR by sterols operate independent of SREBP pathway. INSIG dependent ER-associated protein degradation (ERAD) mechanism is the major post-translational regulatory step mediated by sterol or non-sterol intermediates that involves ubiquitin-proteasomal degradation of HMGCR (6). In view of crucial role of sterols in HMGCR regulation, do intracellular cholesterol levels modulate SHR/WKY-*Hmgcr* promoter activity? Indeed, exogenous cholesterol treatment diminished the SHR/WKY-promoter activity while cholesterol depletion resulted in augmented SHR/WKY-promoter activity (Fig.7A, B). What could be the molecular mechanism of this response to cholesterol level by these promoters? Since Runx3 and Srebf1 are involved in the differential expression of SHR and WKY-*Hmgcr* promoters under basal conditions, we probed their levels upon modulating the intracellular cholesterol level. Interestingly, the Runx3 protein level was also modulated upon cholesterol treatment/depletion (Fig.7E). Consistent with previous reports, the Srebf1 level was enhanced/reduced when cholesterol was diminished/augmented (Fig.7F) (50). Thus, MCD treatment enhanced Runx3 protein level thereby, increasing the Runx3-mediated repression of SHR/WKY-Hmgcr promoter activities. On the other hand, elevated cholesterol level reduced Runx3 protein level and hence, augmented the SHR/WKY-*Hmgcr* promoter activity. In corroboration, cholesterol/MCD treatment caused diminished/enhanced Srebf1-mediated activation of SHR/WKY-Hmgcr promoter activities. These results suggest that cholesterol modulates Srebf1 and Runx3 levels thereby, regulating *Hmgcr* expression.

### Rat models of hypertension show diminished serum cholesterol levels

In line with the enhanced Hmgcr mRNA and protein levels in liver tissues of WKY than SHR (Fig.1), multiple studies reported elevated plasma and hepatic cholesterol levels in WKY as compared to SHR (60, 61). We speculate that the differential expression of Hmgcr, resulting in altered serum cholesterol levels, may be a sex-independent phenomenon since SHR strains show similar fold-diminishment (∼1.4-fold, Fig.S3) in serum cholesterol levels with respect to their sex-matched WKY controls. Thus, SHR emerges as a model where dyslipidemia and hypertension do not seem to be inter-connected. This is not surprising as these animals were derived solely based on high blood pressure as the selection criteria (10, 11) possibly resulting in segregation of genes. This is reminiscent of a previous study that revealed diminished hepatic *Hmgcr* expression and plasma cholesterol level in genetically hypertensive BPH (blood pressure high) mice as compared to hypotensive BPL (blood pressure low) mice (8). Additionally, chromogranin A gene knock out mice which displayed severe hypertension, also showed enhanced Hmgcr mRNA level than their wild-type control mice (62). These studies highlight the possibility that modulated *Hmgcr* expression might be a crucial aspect of hypertension and associated cardiovascular complications in mammals. Indeed, serum cholesterol and systolic blood pressure levels displayed a significant inverse correlation (Fig.S4) across various genetically-inbred hypertensive rat strains. We speculate that the decrease in serum cholesterol levels might be a compensatory response to elevated blood pressure in these genetic models of hypertension thereby, alleviating further risk of cardiovascular complications.

### Conclusion and perspectives

This study identified two functional promoter variations in rat models of human essential hypertension that regulate *Hmgcr* expression by altering the binding affinity of Srebf1 and Runx3 with *Hmgcr* promoter under basal and pathophysiological conditions. Our overall observations are shown by a schematic representation (Fig.8). Notably, the expression pattern of *HMGCR* and *SREBF1* correlated well in various human tissues. However, additional investigations are required to unravel the functional implications of these variations for hypertension. Nonetheless, this is the first report on identification and functional characterization of polymorphisms in the *Hmgcr* locus in hypertensive SHR vs. normotensive WKY. Our findings also reveal that multiple genetic rodent models of hypertension (apart from SHR) exhibit diminished serum cholesterol levels possibly as a protective mechanism to attenuate further cardiovascular complications. Future studies may provide novel insights into molecular links of *Hmgcr* with hypertension and other cardiovascular disease states.

**Fig.8.**
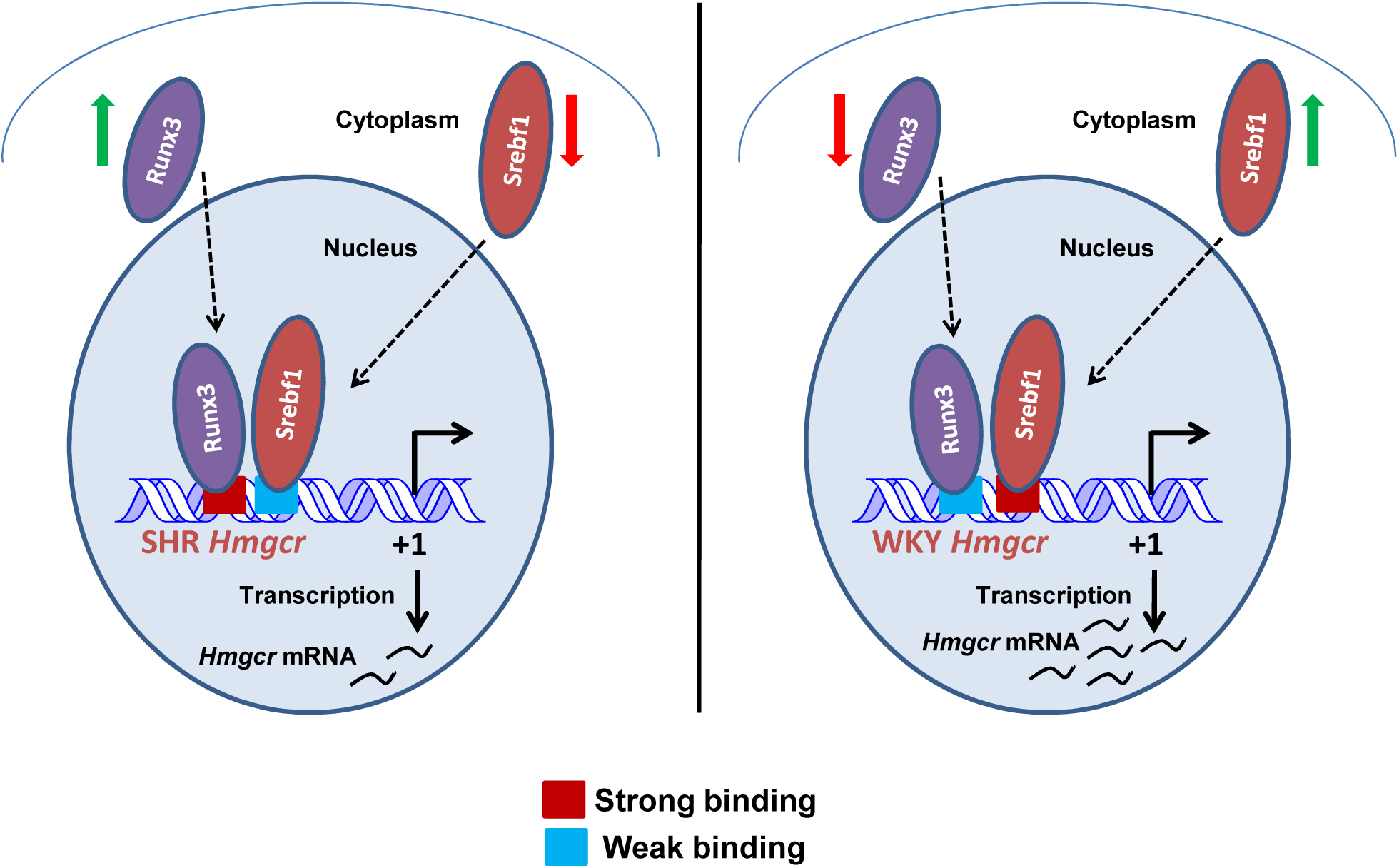
A schematic illustration of mechanism of differential expression of Hmgcr in rat models of hypertension. Runx3 and Srebf1 bind to the SHR-and WKY-*Hmgcr* promoter domain with different affinities (red box: higher affinity, blue box: lower affinity) leading to augmented expression of WKY-*Hmgcr* promoter than the SHR-*Hmgcr* promoter. Moreover, enhanced Runx3 and reduced Srebf1 levels in SHR liver tissues result in lower *Hmgcr* expression.

## ABBREVIATIONS

Hmgcr: 3-Hydroxy-3-Methyl Glutaryl-Coenzyme A reductase;
SHR: spontaneously hypertensive Rat;
WKY: wistar/kyoto; BPH, blood pressure high;
BPL: blood pressure low;
BPN: blood pressure normal;
MCD: methyl-β-cyclodextrin;
ChIP: chromatin immunoprecipitation;
qPCR: quantitative real-time PCR;
RIPA: radio-immunoprecipitation assay;
DMEM: Dulbecco’s Modified Eagle’s Medium;
SREBP-1: Sterol-regulatory element-binding protein 1;
Runx3: runt-related transcription factor 3, PVDF, polyvinylidene difluoride;
ERAD: ER-associated protein degradation

## ACKNOWLEDGEMENTS

This work was supported by Council of Scientific and Industrial Research (CSIR), Government of India to NRM (project number: 37(1564)/12-EMR-II). Research fellowships were received from Ministry of Human Resource Development (to A. A. Khan and V. Gupta), Department of Science and Technology (to V. Arige), Indian Council of Medical Research (to S. S. Reddy) and IIT Madras (to P. Sundar). The authors are grateful to Dr. S-C. Bae (Chungbak National University, South Korea) for providing the pcDNA3.1-Flag-RUNX3 expression plasmid, and to Dr. Hitoshi Shimano (University of Tsukaba, Japan) for the SREBP1a expression plasmid. The authors thank P. Shinde and Dr. T. S. Santra, (Department of Engineering Design, IIT Madras, India) for fluorescence imaging. The authors also thank Dr. S. Lakshmi (IIT Madras, India) and B. Natarajan (IIT Madras, India) for their technical supports to the manuscript.

## AUTHOR CONTRIBUTIONS

A. A. Khan designed, performed experiments, analyzed data and wrote the manuscript. P. Sundar and V. Gupta performed experiments and analyzed data. V. Arige and S. S. Reddy performed experiments. M. Dikshit provided equipment and reagents. M.K. Barthwal provided equipment, reagents and analyzed data. N. R. Mahapatra conceived and designed the study, analyzed data and wrote the manuscript.

