## supplementary information for "Functional promoter polymorphisms govern the differential expression of HMG-CoA Reductase gene in rat models of essential hypertension"

*by*

Abrar A. Khan^1^, Poovitha Sundar^1^, Vinayak Gupta^1^, Vikas Arige^1^, S. Santosh Reddy^2,3^, Madhu Dikshit^4^, Manoj K. Barthwal^2^, Nitish R. Mahapatra^1^

*from the*

^1^Department of Biotechnology, Bhupat and Jyoti Mehta School of Biosciences, Indian Institute of Technology Madras, Chennai 600036, India

^2^Pharmacology Division, CSIR-Central Drug Research Institute, Lucknow 226031, India

^3^Academy of Scientific and Innovative Research (AcSIR), New Delhi 110025, India

^4^Translational Health Science and Technology Institute, Faridabad 121001, India

**Address for correspondence:**

Dr. N. R. Mahapatra, Department of Biotechnology, Bhupat and Jyoti Mehta School of Biosciences, Indian Institute of Technology Madras, Chennai 600036, India. Tel: 91-44-2257-4128;

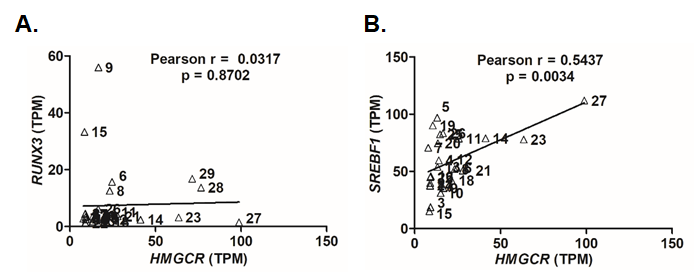

**Fig.S1: Correlation between HMGCR and RUNX3/SREBF1 expression across different human tissues.** Correlation between transcript levels of (A) HMGCR and RUNX3, (B) HMGCR and SREBF1 in various human tissues [n=29; the identity of the tissue samples is given in Table S2] using data retrieved from GTEx Portal (<https://gtexportal.org/home/>). The transcript levels of HMGCR showed a significant positive correlation with the transcript levels of SREBF1 while no correlation was observed between the transcript levels of RUNX3 and HMGCR.

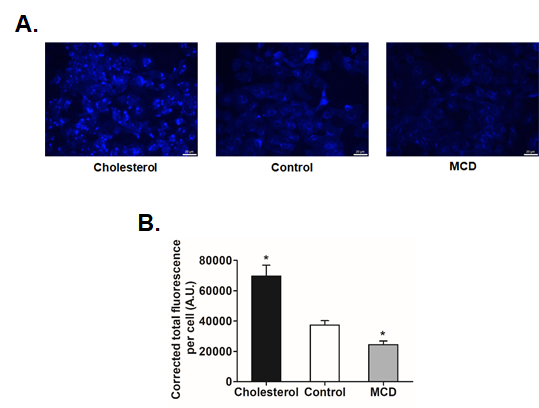

**Fig.S2: Filipin staining of intracellular cholesterol in BRL 3A cells.** (A) The cholesterol/ MCD treatments in BRL 3A cells were confirmed using Filipin stain followed by fluorescence microscopy. Scale bar: 20 µm. (B) Quantification for total corrected fluorescence per cell was performed after cholesterol/MCD treatment using ImageJ. Statistical significance was determined by Student’s *t*-test (unpaired, 2-tailed). *p<0.05 as compared to the control condition.

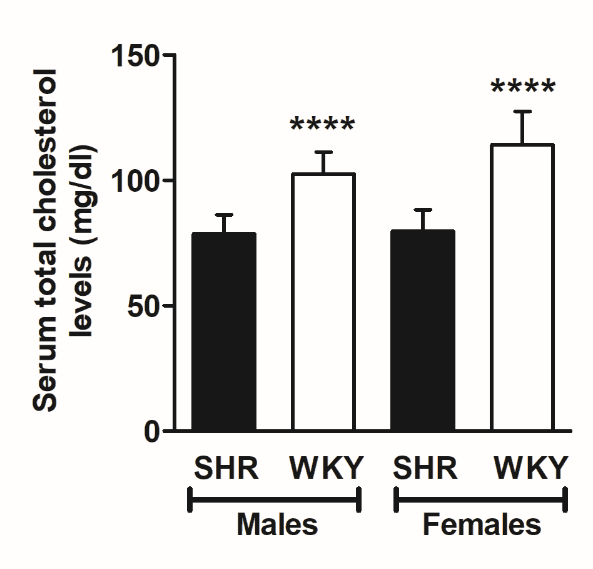

**Fig.S3: Serum total cholesterol levels in SHR and WKY strains.** Serum cholesterol levels for SHR males (n=11), SHR females (n=10), WKY males (n=10) and WKY females (n=14) aged 77-91 days were retrieved from PhenoMiner tool of the Rat Genome Database (<http://rgd.mcw.edu/rgdweb/search/qtls.html?100>). Statistical significance was determined by Student’s *t*-test (unpaired, 2-tailed). ****p<0.0001 with respect to sex-matched SHR group.

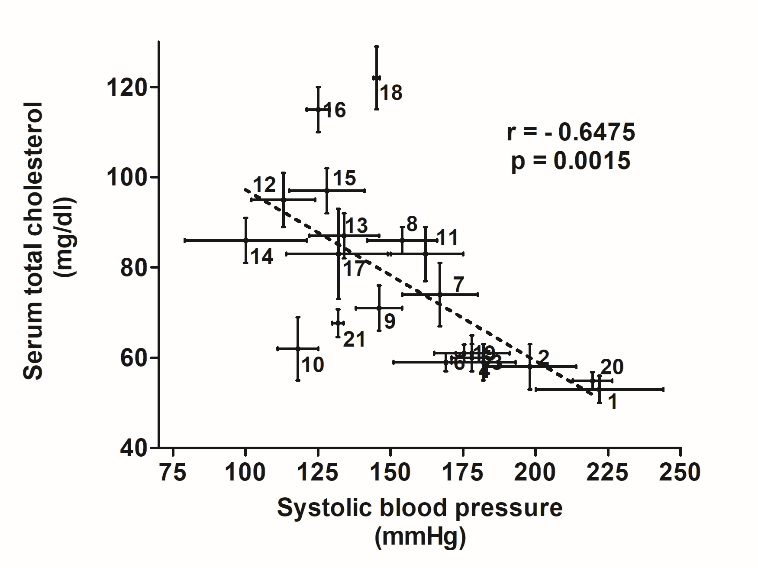

**Fig.S4: Inverse correlation between serum total cholesterol levels and systolic blood pressure in rat models of hypertension.** Correlation between serum cholesterol levels and systolic blood pressure in various genetic rat models of hypertension [n=21; the identity of strains (indicated by numbers) is provided in Table 3] using data from National BioResource Project- Rat (<http://www.anim.med.kyoto-u.ac.jp/nbr/>) and PhenoMiner tool of the Rat Genome Database (<http://rgd.mcw.edu/rgdweb/search/qtls.html?100>).

**Table S1. LOD score and genomic position of various blood pressure QTLs present on rat chromosome 2***

| **Blood Pressure QTLs** | **LOD Score** | **Start position (bp)** | **Stop position (bp)** |
| --- | --- | --- | --- |
| Bp115 | 4.3 | 1 | 35028452 |
| Bp36 | 2.9 | 3125162 | 1.25E+08 |
| Bp240 | 4 | 45317613 | 1.03E+08 |
| Bp243 | 3.9 | 45317613 | 1.55E+08 |
| Bp205 | 3.476 | 45399390 | 62179292 |
| BpQTLCluster2 | 2.86 | 59248567 | 1.25E+08 |
| Bp174 | 4.09 | 61825950 | 79917090 |
| Bp14 | 2.2 | 62179292 | 2.36E+08 |
| Bp203 | 4.377 | 74641110 | 2.24E+08 |
| Bp206 | 3.62454 | 75985115 | 2.36E+08 |
| Bp99 | 2.5 | 84554681 | 1.25E+08 |
| Bp13 | 3.3 | 98037122 | 1.25E+08 |
| Bp6 | 7 | 1.02E+08 | 1.47E+08 |
| Bp296 | 2.1 | 1.03E+08 | 1.48E+08 |
| BpQTLCluster3 | 2.76 | 1.6E+08 | 2.36E+08 |
| Bp357 | 4.1 | 1.63E+08 | 2.08E+08 |
| Bp202 | 3.66819 | 1.67E+08 | 2.59E+08 |
| Bp101 | 2.4 | 1.77E+08 | 2.36E+08 |
| Bp322 | 3.58 | 1.78E+08 | 2.36E+08 |
| Bp295 | 3.5 | 1.81E+08 | 2.26E+08 |
| Bp290 | 1.9 | 2.01E+08 | 2.46E+08 |
| Bp204 | 3.61192 | 2.01E+08 | 2.59E+08 |
| Bp16 | 5.64 | 2.04E+08 | 2.49E+08 |
| Bp63 | 5.64 | 2.04E+08 | 2.49E+08 |
| Bp19 | 6.3 | 2.06E+08 | 2.09E+08 |
| Bp10 | 3.42 | 2.23E+08 | 2.5E+08 |
| Bp158 | 2.62 | 2.23E+08 | 2.5E+08 |
| Bp50 | 4.4 | 2.24E+08 | 2.69E+08 |
| Bp175 | 3.46 | 2.49E+08 | 2.84E+08 |

*This data was retrieved from Rat Genome Database (<http://rgd.mcw.edu/rgdweb/search/qtls.html?100>)

**Table S2. Transcript levels of HMGCR, RUNX3 and SREBF1 across various human tissues mined from GTEx Portal***

| **No.** | **Tissue sample** | ***HMGCR***  **(TPM)** | ***RUNX3***  **(TPM)** | ***SREBF1***  **(TPM)** |
| --- | --- | --- | --- | --- |
| 1 | Adipose-subcutaneous | 9.37 | 3.47 | 45.9 |
| 2 | Adipose-visceral | 9 | 4.5 | 39.83 |
| 3 | Aorta | 9.2 | 3.2 | 18.74 |
| 4 | Bladder | 14.06 | 2.9 | 59.7 |
| 5 | Liver | 13.1 | 1.35 | 96.88 |
| 6 | Lung | 24.64 | 15.67 | 53.11 |
| 7 | Prostate | 7.84 | 2.78 | 70.73 |
| 8 | Small intestine-ileum | 23.26 | 12.51 | 52.06 |
| 9 | Spleen | 16.7 | 55.94 | 35.35 |
| 10 | Stomach | 15.13 | 2.67 | 31.09 |
| 11 | Testis | 25.82 | 5 | 78.65 |
| 12 | Thyroid | 20.25 | 2.365 | 60.45 |
| 13 | Uterus | 13.32 | 1.77 | 53.77 |
| 14 | Vagina | 41.3 | 2.39 | 78.91 |
| 15 | Whole blood | 8.5 | 33.3 | 15.2 |
| 16 | Coronoary artery | 9.4 | 2.41 | 45.38 |
| 17 | Tibial artery | 8.95 | 4.33 | 37.27 |
| 18 | Spinal cord | 21.52 | 1.6 | 42.08 |
| 19 | Ectocervix | 10.58 | 2.55 | 90.14 |
| 20 | Endocervix | 13.52 | 3.7 | 74.58 |
| 21 | Transverse colon | 27.95 | 3.55 | 50.66 |
| 22 | Gastroesophageal Junction | 9.12 | 1.175 | 44.85 |
| 23 | Esophagus-mucosa | 63.52 | 3.15 | 77.87 |
| 24 | Esophagus-muscularis | 8.92 | 1.21 | 37.7 |
| 25 | Fallopian tube | 14.77 | 3.33 | 82.32 |
| 26 | Minor salivary gland | 16.57 | 6.17 | 83.28 |
| 27 | Nerve tibial | 98.83 | 1.44 | 111.9 |
| 28 | Skin-not exposed | 76.62 | 13.7 | 66.16 |
| 29 | Skin-sun exposed | 71.45 | 16.78 | 65.16 |

*This data was retrieved from the GTEx Portal (<https://gtexportal.org/home/>). Tissues with a TPM of at least 1 for *HMGCR*, *SREBF1* and *RUNX3* are shown in the table. TPM: Transcript per million.
